# Fetal programming by the parental microbiome of offspring behavior, and DNA methylation and gene expression within the hippocampus

**DOI:** 10.1101/2024.04.12.589237

**Authors:** Kevin L. Gustafson, Susheel Bhanu Busi, Zachary L. McAdams, Rachael E. McCorkle, Pavlo Khodakivskyi, Nathan J. Bivens, Daniel J. Davis, Murugesan Raju, Lyndon M. Coghill, Elena A. Goun, James Amos-Landgraf, Craig L. Franklin, Paul Wilmes, Rene Cortese, Aaron C. Ericsson

**Affiliations:** Department of Veterinary Pathobiology, College of Veterinary Medicine, University of Missouri, Columbia, MO, 65201, USA; UK Centre for Ecology and Hydrology, Wallingford, Oxfordshire, OX10 8BB, UK; College of Veterinary Medicine, University of Missouri, Columbia, MO, 65211, USA; Department of Chemistry, College of Arts and Science, University of Missouri, Columbia, MO, 65211, USA; University of Missouri Genomics Technology Core, University of Missouri, Columbia, MO, 65211, USA; University of Missouri Bioinformatics and Analytics Core, University of Missouri, Columbia, MO, 65211, USA; Department of Life Sciences and Medicine, Faculty of Science, Technology and Medicine, University of Luxembourg, L-4362 Esch-sur-Alzette, Luxembourg; Department of Child Health & Obstetrics, Gynecology, and Women’s Health, School of Medicine, University of Missouri, Columbia, MO, 65212, USA

**Keywords:** Fetal programming, Maternal microbiome, Gut-brain axis, DNA methylation, Hippocampus, Gene expression, Bile acids, Neurodevelopment

## Abstract

**Background:** The microorganisms colonizing the gastrointestinal tract of animals, collectively referred to as the gut microbiome, affect numerous host behaviors dependent on the central nervous system (CNS). Studies comparing germ-free mice to normally colonized mice have demonstrated influences of the microbiome on anxiety-related behaviors, voluntary activity, and gene expression in the CNS. Additionally, there is epidemiologic evidence supporting an intergenerational influence of the maternal microbiome on neurodevelopment of offspring and behavior later in life. There is limited experimental evidence however directly linking the maternal microbiome to long-term neurodevelopmental outcomes, or knowledge regarding mechanisms responsible for such effects.

**Results:** Here we show that that the maternal microbiome has a dominant influence on several offspring phenotypes including anxiety-related behavior, voluntary activity, and body weight. Adverse outcomes in offspring were associated with features of the maternal microbiome including bile salt hydrolase activity gene expression (*bsh*), abundance of certain bile acids, and hepatic expression of *Slc10a1*. In cross-foster experiments, offspring resembled their birth dam phenotypically, despite faithful colonization in the postnatal period with the surrogate dam microbiome. Genome-wide methylation analysis of hippocampal DNA identified microbiome-associated differences in methylation of 196 loci in total, 176 of which show conserved profiles between mother and offspring. Further, single-cell transcriptional analysis revealed accompanying differences in expression of several differentially methylated genes within certain hippocampal cell clusters, and vascular expression of genes associated with bile acid transport.

Inferred cell-to-cell communication in the hippocampus based on coordinated ligand-receptor expression revealed differences in expression of neuropeptides associated with satiety.

**Conclusions:** Collectively, these data provide proof-of-principle that the maternal gut microbiome has a dominant influence on the neurodevelopment underlying certain offspring behaviors and activities, and selectively affects genome methylation and gene expression in the offspring CNS in conjunction with that neurodevelopment.

## Background

Neurodevelopmental and behavioral disorders are a growing concern worldwide. According to the World Health Organization, nearly one billion people worldwide live with a mental disorder^1^. Behavioral disorders are commonly associated with social impairments, decreased productivity, financial losses, and general maladjustment^2^. Previous studies have found that such disorders contribute substantially to global nonfatal health loss^1^. Anxiety disorders (AD) are increasing in prevalence, affecting close to 1 in 10 children and adolescents between the ages of 3 and 17^3^. Like most mood disorders, AD are multifactorial and often result from a combination of genetic, environmental, and experiential factors. Similarly, one in five children in the U.S. are obese or overweight^4^, reflecting the combined influence of western diet, increasingly sedentary lifestyles, and other factors. Moreover, AD and obesity/overweight (OO) are reciprocal risk factors, frequently occurring as co-morbidities^5–9^.

A growing body of research has linked the gut microbiome (GM) to neurodevelopment and behavior^10–15^, and growth rate or weight gain^16–19^. Work with germ-free mice shows the importance of the GM in normative behavior and metabolism^20–22^, and transfer of an anxiety-related phenotype or increased energy harvest via fecal microbiome transfer indicates that certain features within naturally occurring microbiomes influence these phenotypes^23–25^. There is also evidence that indicates that the effects of the GM can go beyond simply influencing the host. Research in rodents has revealed that the GM of a pregnant dam can influence the fetus and phenotype of the offspring following birth. Eloquent studies in mice have shown that effects of diet and exercise on the maternal GM can be transferred to the offspring, relieving negative metabolic phenotypes ^26, 27^. There are also developmental components to both AD and OO, raising the question of how the maternal microbiome during pregnancy affects fetal development and subsequent behavior and energy metabolism in the adult offspring. The maternal gut microbiome during pregnancy produces metabolites which reach peripheral circulation and the fetal CNS^28^, and maternal proteins and peptides produced by enteroendocrine cells in response to the microbiome likely also cross the placenta and reach fetal circulation^29–31^. Disruption of the maternal GM can affect these processes as demonstrated by increased anxiety in the offspring of mice with antibiotic- or diet-induced dysbiosis^32–35^.

There are still major gaps in our knowledge however regarding the mechanisms through which the maternal microbiome during pregnancy programs long-lasting changes in offspring behavior and metabolism. These intergenerational effects suggest fetal imprinting by an unknown mechanism, while differences in anxiety-related behavior (and other complex behaviors) indicate a neurodevelopmental basis. Owing to the genetic, dietary, and environmental heterogeneity, analysis of these processes in a human population requires very large sample sizes and long-term tracking of mother-child pairs. To circumvent these factors, here we use two groups of genotype-, age-, and sex-matched outbred CD-1 mice consuming the same diet. To be clear, all mice in these two colonies are of the same genetic background, and only differ in the two microbiomes they harbor. These microbiomes, originally derived from Jackson Laboratory and Envigo (now known as Inotiv), are characterized by low and high alpha diversity relative to each other and distinct beta diversity. These two colonies were developed at MU Mutant Mouse Resource and Research Center (MMRRC) by initially transferring CD-1 embryos into respective C57BL/6 dams and allowing the dams to transfer their GMs to offspring via natural postnatal transmission. These CD-1 pups became the founders of these two colonies which have been maintained and continually monitored for GM stability within our facility for over 35 generations. Additionally, a rotational breeding scheme and routine introduction of CD-1 genetics via embryo transfer from CD-1 mice purchased from Charles River allows for the maintenance of allelic heterozygosity within each colony and ensures these colonies do not become genetically distinct from each other. Since CD-1 mice that harbor a Jackson Laboratory origin GM have a GM with low phylogenetic richness and diversity, the GM of these mice was designated GM^Low^. The CD-1 colony with an Envigo origin GM has relatively high phyologenetic diversity and is thus designated GM^High^. Phenotypic assessments of these two colonies revealed differences in anxiety-related behavior, voluntary activity, fetal growth, food intake, and adult growth^36, 37^.

We hypothesized that the maternal GM would influence the neurodevelopment of the offspring via fetal programming while *in utero* by GM-derived metabolites. Taking advantage of the phenotypic differences in these two colonies, we utilized cross-foster studies to determine the relative influence of the parental (i.e., prenatal) and offspring (i.e., postnatal) microbiome on offspring phenotypes. Cross-fostering between dams of the reciprocal GM (e.g., pups born to GM^Low^ cross-fostered to GM^High^, and vice versa) allows offspring to develop under the influence of the birth dam GM *in utero*, and then acquire the surrogate dam GM during postnatal life. Microbiome-associated differential phenotypes in which cross-fostered offspring match the phenotypes observed in surrogate dams suggest a postnatal influence, while similarities between cross-fostered offspring and their birth dams suggest a dominant prenatal influence of the parental microbiome. Here, we expand on previous behavioral phenotyping to include control and cross-fostered (CF) offspring, demonstrating a dominant influence of the birth dam GM on offspring development and behavior at seven weeks of age. This work was complemented by microbial and metabolic profiling of mice in each colony, genome-wide methylome analysis of hippocampal DNA from dams and control and CF offspring, and single nuclei transcriptome analysis of RNA from control and CF offspring. Previously identified phylogenetic differences are now complemented by differences in certain metabolites, including bile acids (BA), and differential ileal and hepatic expression of BA receptors and transporters. Analysis of hippocampal DNA revealed dominant effects of the maternal microbiome on CpG methylation, maintained in offspring independent of the postnatal microbiome. Single-nuclei RNA sequencing (snRNA-seq) of hippocampal RNA confirmed fetal programming of several cell-specific differentially methylated genes, including genes involved in G protein-coupled receptor and orexigenic signaling pathways.

## Methods

### Mice

All mice tested in the current study were outbred CD-1 mice (Crl:CD1(ICR)) generated from breeders obtained from the Mutant Mouse Resource and Research Center at the University of Missouri (MU MMRRC). Multiple different cohorts of mice were utilized for various outcomes. CD-1 mice were from two colonies in which the founders were originally purchased from Charles River (Frederick, MD), and were generated via rederivation to harbor either a high richness Envigo (now Inotiv, Indianapolis, IN) origin GM (GM^High^), or a low richness Jackson Laboratory origin GM (GM^Low^) as previously described^24^. All donor mice were reared at the MU MMRRC and the two colonies have been maintained and continually monitored for GM stability within our facility for over 35 generations. Additionally, a rotational breeding scheme and routine introduction of CD-1 genetics via embryo transfer from CD-1 mice purchased from Charles River allows for the maintenance of allelic heterozygosity within each colony and ensures these colonies do not become genetically distinct from each other. Since CD-1 mice that harbor a Jackson Laboratory-origin GM were found to have a GM with low phylogenetic richness and diversity, the GM of these mice was designated GM^Low^. Similarly, since CD-1 mice that harbored an Envigo-origin GM were found to have a GM with high phylogenetic richness and diversity relative to GM^Low^, the GM of these mice was designated GM^High^. Colonies of mice were housed under barrier conditions in microisolator cages with compressed pelleted paper bedding and nestlets, on ventilated racks with *ad libitum* access to irradiated chow and acidified, autoclaved water, under a 14:10 light/dark cycle. Mice were determined to be free of all bacterial pathogens including *Bordetella bronchiseptica*, *Filobacterium rodentium*, *Citrobacter rodentium, Clostridium piliforme, Corynebacterium bovis, Corynebacterium kutscheri, Helicobacter* spp., *Mycoplasma* spp., *Rodentibacter* spp., *Pneumocystis carinii, Salmonella* spp., *Streptobacillus moniliformis, Streptococcus pneumoniae*; adventitious viruses including H1, Hantaan, KRV, LCMV, MAD1, MNV, PVM, RCV/SDAV, REO3, RMV, RPV, RTV, and Sendai viruses; intestinal protozoa including *Spironucleus muris, Giardia muris, Entamoeba muris,* trichomonads, and other intestinal flagellates and amoebae; intestinal parasites including helminths; and external parasites including all species of lice and mites, via quarterly sentinel testing performed by IDEXX BioAnalytics (Columbia, MO). Fecal samples were collected from pregnant dams at 19 days of gestation, and from mouse pups at time of weaning (21 days of age) using previously described methods^38^. Briefly, mice were placed in an empty autoclaved cage within a biological safety cabinet and allowed to defecate. Freshly evacuated samples feces were immediately collected into a sterile collection tube using autoclaved wooden toothpicks discarded after each single usage. All samples were promptly placed on ice. Following fecal sample collection, samples were stored in a -80°C freezer until DNA extraction was performed. Samples were collected from all experimental mice at 50 days of age, at time of necropsy. All dams were mated with sires of the same GM and were housed together until approximately day 14 of gestation, at which time sires were removed. All dams were singly housed for the last week of gestation to ensure pups were correctly assigned to their birth dams. All dams were handled minimally during gestation, and only handled for routine cage changes by vivarium care staff and once for pre-parturition fecal sample collection. During the one week of anxiety-related behavior testing, only the investigators handled and entered the home cages to avoid unknown and excessive disturbances to the mice.

### Gut microbiome analysis

*DNA extraction*. Fecal DNA was extracted using QIAamp PowerFecal Pro DNA kits (Qiagen), according to the manufacturer’s instructions, with the exception that the initial sample disaggregation was performed using a TissueLyser II (Qiagen), rather than a vortex and adaptor as described in the protocol.

*16S rRNA amplicon library preparation and sequencing*. Extracted fecal DNA was processed at the University of Missouri DNA Core Facility. Bacterial 16S rRNA amplicons were constructed via amplification of the V4 region of the 16S rRNA gene using previously developed universal primers (U515F/806R), flanked by Illumina standard adapter sequences^39, 40^. Oligonucleotide sequences are available at proBase^41^. Dual-indexed forward and reverse primers were used in all reactions. PCR was performed in 50 µL reactions containing 100 ng metagenomic DNA, primers (0.2 µM each), dNTPs (200 µM each), and Phusion high-fidelity DNA polymerase (1U, Thermo Fisher). Amplification parameters were 98°C^(3^ ^min)^ + [98°C^(15^ ^sec)^ + 50°C^(30^ ^sec)^ + 72°C^(30^ ^sec)^] × 25 cycles + 72°C^(7^ ^min)^. Amplicon pools (5 µL/reaction) were combined, thoroughly mixed, and then purified by addition of Axygen Axyprep MagPCR clean-up beads to an equal volume of 50 µL of amplicons and incubated for 15 minutes at room temperature. Products were washed multiple times with 80% ethanol and the dried pellet was resuspended in 32.5 µL EB buffer (Qiagen), incubated for two minutes at room temperature, and then placed on a magnetic stand for five minutes. The final amplicon pool was evaluated using the Advanced Analytical Fragment Analyzer automated electrophoresis system, quantified using quant-iT HS dsDNA reagent kits, and diluted according to Illumina’s standard protocol for sequencing on the MiSeq instrument.

*Bioinformatics*. DNA sequences were assembled and annotated at the MU Informatics Research Core Facility. Primers were designed to match the 5’ ends of the forward and reverse reads. Cutadapt^42^ (version 2.6) was used to remove the primer from the 5’ end of the forward read. If found, the reverse complement of the primer to the reverse read was then removed from the forward read as were all bases downstream. Thus, a forward read could be trimmed at both ends if the insert was shorter than the amplicon length. The same approach was used on the reverse read, but with the primers in the opposite roles. Read pairs were rejected if one read or the other did not match a 5’ primer, and an error-rate of 0.1 was allowed. Two passes were made over each read to ensure removal of the second primer. A minimal overlap of three bp with the 3’ end of the primer sequence was required for removal. The QIIME2^43^ DADA2^44^ plugin (version 1.10.0) was used to denoise, de-replicate, and count ASVs (amplicon sequence variants), incorporating the following parameters: 1) forward and reverse reads were truncated to 150 bases, 2) forward and reverse reads with number of expected errors higher than 2.0 were discarded, and 3) Chimeras were detected using the "consensus" method and removed. R version 3.5.1 and Biom version 2.1.7 were used in QIIME2. Taxonomies were assigned to final sequences using the Silva.v132^45^ database, using the classify-sklearn procedure. The cladogram was constructed with GraPhlAn using genus-level taxonomic classifications^46^. Branch color depicts phylum-level classification. The outer ring denotes Benjamini-Hochberg-corrected *p* values from Wilcox Rank-Sum tests comparing the relative abundance of each genus between GMs. The color of the outer ring indicates the GM with the greater average relative abundance of that genus.

### Real-time reverse transcription-polymerase chain reaction (qRT-PCR)

Total RNA was isolated from tissues using the Qiagen RNeasy Mini Kit per manufacturer’s instructions. RT-PCR was performed using the BioRad iTaq Universal SYBR Green One-Step Kit following the manufacturer instructions. Briefly, each reaction consisted of 5 μL of SYBR Green Supermix, 0.125 μL iScript reverse transcriptase, 0.45 μL of forward and reverse primers, 1.475 μL of water, and 2.5 μL of template RNA. The reaction was run on a BioRad C1000 Touch thermal cycler with a BioRad CFX384 Real-Time System with the following parameters: 50°C for 10 min for reverse transcription, 95°C for 1 min for DNA Polymerase activation and DNA denaturation, and 40 cycles of 95°C for 10 sec and 60°C for 30 sec. Melt-curve analysis was performed using the following parameters: 65-95°C with 0.5°C increments for 5 sec/step. Primers used can be found in **Table S1**.

### Bile salt hydrolase metagenomic and metatranscriptomic analysis of mouse feces

Metagenomic and metatranscriptomic *bsh* (K01442) read counts were acquired from previous multi-omic analysis of GM^Low^ (GM1) and GM^High^ (GM4)^47^. Expression counts from metatranscriptomic analysis were normalized to *bsh* metagenomic reads.

### Measurement of BSH activity in mouse feces

BSH activity in the mouse feces was measured using previously reported bioluminescent bile acid activatable luciferin probes (BAL) protocol^48^ with major modification by replacing whole-cell bioluminescence readout with the recombinant luciferase enzymatic assay^49^. Mouse fecal samples were soaked in PBS (pH 7.4, Gibco, ref# 10010-023) supplemented with 2- mercaptoethanol (Acros Organics, 20 mM) at a concentration of 10 mg/mL on ice for 30 min. The mixtures were homogenized by sonication in ultrasound cleaner (Elmasonic Easy 40 H, 340 W) at 0°C for 30 min, stirring every 10 min. Resulting mixtures along with blank buffer (3 replicates by 50 μL) were mixed with working solutions of BAL probes (50 μL, 20 μM in PBS) along with a solution of luciferin (50 μL, 2 μM in PBS) in a 96-well assay plate (Corning, ref# 3595) and incubated at 37°C for 1 h. After incubation, the mixtures were diluted with 2% Triton X-100 in PBS (100 μL) to stop the reaction. In a separate 96-well flat bottom black plate (Corning, ref# 3650), resulting mixtures (5 μL) were diluted with PBS (50 μ). A luciferase solution containing recombinant luciferase from Photinus pyralis (Sigma-Aldrich, 20 μg/mL), ATP disodium trihydrate (Fisher Scientific, 2 mM), and magnesium sulfate heptahydrate (Fisher Scientific, 2 mM) in PBS (50 μL) was added to each well simultaneously. Bioluminescence was measured immediately in an IVIS Spectrum (Xenogen) imaging system for 20 min with 1 min intervals using automatic settings. Raw data were processed using Living Image 4.2 software (Caliper LifeSciences), further data processing was carried out in Excel (Microsoft 365), and finally visualization and statistical calculations were performed in Prism 9 (GraphPad software). Deconjugation potentials or percentage of probe hydrolysis were calculated as the ratio of the signal from the BAL probe to the signal from luciferin in the corresponding fecal extract and reported as the mean value of 3 replicates. The signals from incubation of BAL probes in blank buffer provided a background result of nonspecific hydrolysis of the probes.

### Metabolite analyses

*GC-MS.* Fecal and serum samples were diluted in 18 volumes of ice-cold 2:2:1 methanol/acetonitrile/water containing a mixture of internal standards (D4-citric acid, D4- succinic acid, D8-valine, and U13C-labeled glutamine, glutamic acid, lysine, methionine, serine, and tryptophan; Cambridge Isotope Laboratories), where the 1-part water was composed of sample volume + water. Sample extraction mixtures were vortexed for 10 minutes at RT and rotated for 1 hour at −20°C. Mixtures were centrifuged for 10 minutes at 21,000 × g, and 150 µL of the cleared metabolite extracts were transferred to autosampler vials and dried using a SpeedVac vacuum concentrator (Thermo). Dried metabolite extracts were reconstituted in 30 μL of 11.4 mg/mL methoxyamine (MOX) in anhydrous pyridine, vortexed for 5 minutes, and heated for 1 hour at 60°C. Next, to each sample 20 μL of N,O-Bis(trimethylsilyl)trifluoroacetamide (TMS) was added, samples were vortexed for 1 minute, and heated for 30 minutes at 60°C. Derivatized samples were analyzed by GC-MS. One μL of derivatized sample was injected into a Trace 1300 GC (Thermo) fitted with a TraceGold TG-5SilMS column (Thermo) operating under the following conditions: split ratio = 20:1, split flow = 24 μL/minute, purge flow = 5 mL/minute, carrier mode = Constant Flow, and carrier flow rate = 1.2 mL/minute. The GC oven temperature gradient was as follows: 80°C for 3 minutes, increasing at a rate of 20°C/minute to 280°C, and holding at a temperature at 280°C for 8 minutes. Ion detection was performed by an ISQ 7000 mass spectrometer (Thermo) operated from 3.90 to 21.00 minutes in EI mode (-70eV) using select ion monitoring (SIM).

*LC-MS SCFA analysis.* 18-fold (w/v) extraction solvent (Acetonitrile:Methanol:Water (2:2:1)) containing deuterated SCFA standards (D3-acetate, D7-butyrate, and D5-propionate) was added to each sample and rotated at −20°C for 1 hr and then centrifuged at 21,000 × g for 10 min. Supernatant was used for LC-MS SCFA analysis. LC-MS data was acquired on a Thermo Q Exactive hybrid quadrupole Orbitrap mass spectrometer with a Vanquish Flex UHPLC system or Vanquish Horizon UHPLC system. The LC column used was a ZIC-pHILIC guard column (20 × 2.1 mm). The injection volume was 2 µL. For the Mobile phase, Solvent A consisted of 20 mM ammonium carbonate [(NH4)2CO3] and 0.1% ammonium hydroxide (v/v) [NH4OH] at pH ∼9.1] and Solvent B consisted of Acetonitrile. This method was run at a flow rate of 0.1 mL/min, and the injection volume was 2 µL. Linear gradient was used at 70% solvent B with a 5 min elution time. The mass spectrometer was operated in targeted selected ion-monitoring (tSIM) mode from 1 to 5 minutes. An inclusion list for the three short chain fatty acids and their deuterated versions were used in tSIM method.

*LC-MS bile acid analysis.* Extraction solvent (methanol:acetonitrile:water, 2:2:1) was spiked with (5 µLl/mL) deuterated bile acids MaxSpec Mixture (Cayman Chemicals Item no. 33506). 18-fold volume extraction buffer was added to each sample. The samples were placed in a −20°C freezer for 1 hour while rotating. The samples were then centrifuged at 21,000 × g for 10 minutes. Supernatant was transferred to LC-MS autosampler vials for analysis. LC-MS data was acquired on a Thermo Q Exactive hybrid quadrupole Orbitrap mass spectrometer with a Vanquish Flex UHPLC system or Vanquish Horizon UHPLC system. A Thermo Hypersil GOLD (2.1 × 150 mm, 1.9 µm) UHPLC column was used with a column Temperature of 30°C. For the Mobile Phase, solvent A consisted of 1% acetonitrile in water with 0.1% formic acid, and solvent B is 99% acetonitrile with 0.1% formic acid. The gradient started at 50% Solvent B and was held for 2.5 minutes; then increased to 100% B at 10 minutes and held for 0.5 minutes before re-equilibration to 50% solvent B for 5.5 min. Flow Rate was 0.4 mL/min, and injection volume was 3 µL. The mass spectrometer was operated in full-scan negative mode, with the spray voltage set to 3.0 kV, the heated capillary held at 275°C, and the HESI probe held at 350°C. The sheath gas flow was set to 40 units, the auxiliary gas flow was set to 15 units, and the sweep gas flow was set to 1 unit. MS data acquisition was performed in a range of m/z 70–1,000, with the resolution set at 70,000, the AGC target at 1 × 10^6^, and the maximum injection time at 200 ms^50^.

*Metabolomic Data Analysis.* GC-MS Raw data were analyzed using TraceFinder 5.1 (Thermo). Metabolite identification and annotation required at least two ions (target + confirming) and a unique retention time that corresponded to the ions and retention time of a reference standard previously determined in-house. A pooled-sample generated prior to derivatization was analyzed at the beginning, at a set interval during, and the end the analytical run to correct peak intensities using the NOREVA tool^51^. NOREVA corrected data were then normalized to the total signal per sample to control for extraction, derivatization, and/or loading effects. Acquired LC-MS data were processed by Thermo Scientific TraceFinder 4.1 software, and metabolites were identified based on the University of Iowa Metabolomics Core facility standard-confirmed, inhouse library. NOREVA was used for signal drift correction^51^. For bile acids, data were normalized to one of the d4-bile acid standards. For SCFA, analyte signal was corrected by normalizing to the deuterated analyte signal and the signal obtained from processing blank was subtracted.

### Cross-fostering

Mice in cross-foster (CF) groups were cross-fostered at less than 24 hours of age to a surrogate dam of the reciprocal GM. Following identification of recently birthed litters from both GMs, cages were moved to a biosafety cabinet. Litters were removed from the cage of the biological dam and placed onto clean paper towels. Bedding from the cage of the surrogate dam was gently mixed with the pups to transfer the surrogate dam scent to the pups and reduce the possibility of cannibalism. The pups were then placed into the surrogate dam cage, and cages were returned to the appropriate housing rack.

### Behavior testing

*Open Field Exploration.* The open field exploration test was used to evaluate anxiety-related behavior and locomotor function. Environmental control chambers (Omnitech Electronics, Inc., Columbus, OH) consisting of 4 separate environmental isolation chambers containing a plexiglass box (41cm × 41cm × 30cm) placed onto an infrared grid (41cm × 41cm) to track locomotion. Lighting for each isolation chamber was set to 159 lux. Mice were allowed to acclimate to the behavior room for 1 hour prior to testing. Before starting each test, the plexiglass was cleaned with 0.25% bleach, followed by 70% ethanol to remove any residual olfactory cues. Each mouse was placed into the middle of the open field exploration test and recorded for 30 minutes by the Fusion behavior monitoring software (Omnitech Electronics, Inc., Columbus, OH). The first 20 minutes of the test were considered acclimation time, and the final 10 minutes were analyzed following completion of the test. Total distance traveled (cm), time spent in the center zone (seconds), distance traveled in the center zone, and vertical activity (rearing) were measured.

*Light/Dark Transition.* The light/dark transition test was performed within the environmental control chambers (Omnitech Electronics, Inc., Columbus, OH, USA). The apparatus consisted of a plexiglass box within the environmental chambers (41cm × 41cm × 30cm) that were partitioned into two equal sections by a black plexiglass insert with a door that allowed one half to be a dark zone, and a second half to be a light zone. The light zone was illumination was set to 200 lux. Mice were allowed to acclimate to the behavior room for 1 hour prior to testing. Before the start of each test, the plexiglass box and insert were cleaned with 0.25% bleach, followed by 70% ethanol to remove any residual olfactory cues. Mice were placed into the light zone facing away from the dark zone and monitored by Fusion behavior monitoring software (Omnitech Electronics, Inc., Columbus, OH, USA) for 15 minutes. Time spent in the light zone (sec), distance travelled in the light zone (cm), and number of transitions between light and dark zones were measured.

*Elevated Plus Maze.* The elevated plus maze test consisted of an apparatus with two open arms (32.5 × 5 cm, with 2-mm ledges) and two closed arms (32.5 × 5 cm, with 14.5 cm high walls). The open arms were arranged perpendicular to the closed arms so the apparatus formed the shape of a plus sign with a center square (5 × 5 cm). The entire apparatus was raised 50 cm above the floor. The center zone of the apparatus was illuminated to 50 lux. Mice were allowed to acclimate to the behavior room for 1 hour prior to testing. Prior to testing, the apparatus was cleaned with 0.25% bleach followed by 70% ethanol to remove olfactory cues. Each mouse was placed in the center square facing an open arm and was recorded and monitored for 5 minutes. Distance Traveled in the open arms (cm), time spent in the open arms (sec), and number of entries into the open arms was calculated from distance measurements and entry counts obtained by Any-Maze monitoring software (Stoelting Co., Wood Dale, IL, USA).

*Voluntary running*. New litters of CD-1 GM^Low^ and GM^High^ mice were generated to evaluate voluntary wheel running assays (the mice used in the behavior assays did not undergo wheel running evaluation). Litters for running wheel experiments were culled to six pups per litter (3 male, 3 female) at birth, and then weaned into cages of same-sex trios at weaning. During wheel set-up at seven weeks of age, mice were transferred from their home cage to a new static microisolator cage containing a wireless running wheel (Med Associates, ENV-047) connected to a wireless hub and laptop computer in the animal room. Only investigators entered the behavior room to check mice and equipment daily during the 12 days of testing to avoid excessive disturbances to the mice. Mice were singly housed during the experiment, assigned to running wheel cages using a random number generator, and were placed in alternating order on the shelf such that microbiome group and sex were consistently alternated. Following five days of acclimation, data were collected continuously for seven consecutive days using Wheel Manager software, v2.04.00 (Med Associates, SOF-860). Data were analyzed using Wheel Analysis software, v2.02.01 (Med Associates, SOF-861). No other mice were housed in the room containing running wheel cages, traffic was limited to once daily checks at the same time of day by one laboratory staff, and no cage changes were performed during the acclimation and testing period.

### Necropsy

Mice were transported to the necropsy room at 50 days of age and allowed to acclimate to the room for 1 hour. Mice were then euthanized one at a time by CO_2_ asphyxiation out of sight of other mice. The euthanasia chamber was cleaned with 70% ethanol between mice to eliminate olfactory cues. Following loss of paw pinch and righting reflexes, blood was collected by cardiac puncture and placed into serum separator tubes. The brain was then removed, and the hippocampus was gently dissected out, placed in a 2 mL tube, and promptly plunged into liquid nitrogen to flash freeze. Liver and ileum were isolated, placed in a 2 mL tube, and promptly plunged into liquid nitrogen. Two fecal pellets were collected from the colon for 16S rRNA amplicon sequencing. Blood was allowed to clot for 30 minutes at room temperature and was then centrifuged at 4,000 RPM for 15 minutes, and serum placed into a 1.5 mL microcentrifuge tube. Hippocampus, feces, and serum were promptly placed into -80°C freezer for storage.

### Methylome analysis

Due to constraints on resources, methylome analysis were performed in only one sex. To assess intergenerational effects on DNA methylation, we analyzed dams and female offspring in both control and CF mice. Dams from both colonies were time mated, and following birth, litters were culled to six female pups. Three of the pups from each litter remained with the birth dam, and three were cross-fostered onto a dam of the opposite GM so that every dam from both GMs had three of their birth pups and three cross-foster surrogate pups. Hippocampi were collected from all dams following weaning, flash frozen in liquid nitrogen, and stored at -80°C. At seven weeks of age, hippocampi from the offspring were collected, flash frozen in liquid nitrogen, and stored at -80°C. Hippocampal genomic DNA was isolated from adult female CF and control offspring hippocampi using the DNeasy kit (Qiagen, Valencia, CA) following manufacturer instructions. For studying genome-wide DNA methylation profiles, 1 μg of genomic DNA was treated with sodium bisulfite (Zymo Research, Irvine, CA). Converted DNA was analyzed using Infinium Mouse Methylation BeadChip assay (Illumina, San Diego, CA). This array includes over 285,000 CpG sites covering all RefSeq genes, including CpG islands, translation start sites, enhancers, imprinted loci, and other regions^52^. All data analyses were conducted using the R environment version 4.2.0. Microarray data was processed using the *ENmix* version.1.34.02^53^ and *minfi* v.1.44.0^54^ packages. Quantile normalization of U or M intensities for Infinium I or II probes were performed, respectively. A model-based correction was performed, and a dye-bias correction was conducted using *RELIC* ^55^. β-values representing the averaged total intensity value per CG position was calculated as unmethylated intensity (U) + methylated intensity (M) [M / (U + M + 100)]. Probes with a detection p < 1 × 10^-6^ and less than 3 beads were defined as low quality. Samples with low quality methylation measurements > 5% or low intensity bisulfite conversion probes were removed from further analysis. Differentially methylated regions (DMRs) between the experimental groups were determined using the *ENmix* version.1.34.02^53^ package. For each position, the magnitude of the DNA methylation difference was expressed as Fold Changes in the logarithmic scale (logFC) and the significance of the difference as a FDR-corrected p value (q value).

### Isolation of hippocampal nuclei

Single nuclei were isolated from mouse hippocampal tissue samples as follows. Briefly, Nuclei Lysis Buffer was prepared by adding 12 mL of Nuclei EZ Prep Lysis Buffer (Sigma-Aldrich, St. Louis, MO, USA) to a 15mL tube and adding 1 cOmplete Ultra tablet (Sigma-Aldrich, St. Louis, MO, USA) and allowing tablet to dissolve. Two 4-mL aliquots of the Nuclei EZ Prep Lysis Buffer + cOmplete Ultra tablets were then placed in 15 mL tubes. Twenty (20) µL of Protector RNase inhibitor (Sigma-Aldrich, St. Louis, MO, USA) and 20 uL of Superase-In (MilliporeSigma, Burlington, MA, USA) were added to one 4 mL aliquot to make Nuclei Lysis Buffer 1 (NLB1). Four (4) µL of Protector RNase inhibitor (Sigma-Aldrich, St. Louis, MO, USA) and 4 µL of Superase-In (MilliporeSigma, Burlington, MA, USA) were added to the second -mL aliquot to make Nuclei Lysis Buffer 2 (NLB2). Suspension Buffer (SB) was prepared by adding 1 mL of fetal bovine serum (Sigma-Aldrich, St. Lous, MO, USA) to 9 mL of 1× phosphate-buffered saline with 4 µL of Protector RNase inhibitor (Sigma-Aldrich, St. Louis, MO, USA). Eight (8) hippocampi halves from eight individual mice were pooled and homogenized to a single cell suspension in a 2 mL dounce homogenizer with 2 mL of NBL1. The sample was then strained through a 200 µm strainer (pluriSelect Life Science, El Cajon, CA, USA) and the strained cell suspension returned to the 2 mL dounce and homogenized to a nuclei suspension. The nuclei were strained through a 40 µm strainer (pluriSelect Life Science, El Cajon, CA, USA) and centrifuged at 500 RCF at 4°C for 5 minutes. Supernatant was removed, and pellet was resuspended with NLB2 and incubated at 4°C for 5 minutes. Nuclei were then centrifuged at 500 RCF at 4°C for 5 minutes, supernatant was removed, and nuclei were resuspended in suspension buffer.

### 10x Genomics single cell 3’ RNA-Seq library preparation

Libraries were constructed by following the manufacturer’s protocol with reagents supplied in 10x Genomics Chromium Next GEM Single Cell 3′ Kit v3.1. Briefly, nuclei suspension concentration was measured with an Invitrogen Countess II automated cell counter. Nuclei suspension (1,200 nuclei per microliter), reverse transcription master mix, and partitioning oil were loaded on a Chromium Next GEM G chip with a capture target of 8,000 nuclei per library. Post-Chromium controller GEMs were transferred to a PCR strip tube and reverse transcription performed on an Applied Biosystems Veriti thermal cycler at 53°C for 45 minutes. cDNA was amplified for 13 cycles and purified using Axygen AxyPrep MagPCR Clean-up beads. Fragmentation of cDNA, end-repair, A-tailing and ligation of sequencing adaptors was performed according to manufacturer specifications. The final library was quantified with the Qubit HS DNA kit and the fragment size determined using an Agilent Fragment Analyzer system. Libraries were pooled and sequenced on an Illumina NovaSeq 6000 to generate 50,000 reads per nuclei.

### Single cell bioinformatics

FASTQ files were generated from the raw base call outputs using Cell Ranger (10x Genomics) pipeline, *mkfastq* v3.0. Using default parameters, a UMI (gene-barcode) count matrix per sample was obtained using the built-in Cell Ranger count pipeline. To reduce noise, we only kept genes that were detected in at least three barcodes, and subsequently removed ribosomal- encoded genes from the count matrix. Scrublet^56^ was then used to identify potential multiplet- barcodes and only those with a doublet score of less than 0.15 were used for downstream analyses. The files were then combined in a single embedding for the control and CF groups separately, following the Seurat v3 integration workflow^57^. SCTransform was used to normalize each sample, followed by the identification of integration anchors and variable features using the Seurat workflow. Dimension reduction was performed scaled data after 4000 highly variable genes across the samples were identified (*SelectIntegrationFeatures* function). The *IntegrateData* function was then used to obtain a combined and centered matrix, where the top 30 components were used to carry out ordination analyses. These components were used to build a SNN (shared nearest neighbour) graph which was subsequently clustered using the *Louvain* algorithm for speed and computational efficiency. The principal components were then mapped into two dimensions using the default uniform manifold approximation and projection (UMAP) algorithm, where the n = 30 neighbours was set, with a minimum distance of 0.4. Finally, the *FindAllMarkers* function was used to identify marker genes for each cluster. The top marker genes were used manually based on literature searches to assign cell type annotations for each cluster. This was further corroborated by cluster annotations using the Azimuth mouse reference datasets^57^. The cell types across samples and groups were combined with their pseudo bulk profiles, and the resulting gene-cell type matrix was normalized by estimating transcripts per million and transformed (log2) for downstream analyses. To obtain statistically enriched differential gene expression, we used generalized additive regression models, where in the control or CF variables, alongside the GM^Low^ or GM^High^ status were encoded as independent variables. The models were analyzed for each cluster independent of the other, where per gene log2 fold-change was determined. Significance was identified as those genes with an adjusted p value of less than 0.05, following Benjamini-Hochberg correction. All figures and statistical analyses were performed using R v4.1^58^.

Cell-to-cell communication was inferred using log10-transformed gene expression data collected from snRNA-seq of the mouse hippocampus using CellChat^59, 60^. Cell clusters as identified above were grouped into the following cell types based on Azimuth classification: glutamatergic and GABAergic neurons, periendothelial cells, microglia, astrocytes, and oligodendrocytes. Ligand-receptor interactions were inferred using the mouse reference database provided by CellChat (Accessed May 25, 2023). Overexpressed genes and interactions were determined using default settings. Cell-cell communication probability was inferred using default settings. Information flow was determined by the summation of communication probability for each pathway.

### Statistics

Two-way Analysis of Variance (ANOVA) followed by Tukey’s *post hoc* analysis was used to test for main effects of GM and sex in OFE, LDT, and EPM behavior tests for all behavior testing parameters. Due to lack of normality, CF LDT parameter distance travelled in the light zone and CF EPM parameters time spent in open arms and distance travelled in open arms were normalized by square root transformation. Two-way ANOVA followed by Tukey’s *post hoc* was used to test for main effects of GM and Time (day) for voluntary running distance data. Two-way ANOVA followed by Tukey’s *post hoc* was used to test for main effects of GM and sex in the body weight data. Two-way Permutational analysis of variance (PERMANOVA) was used to test group beta-diversity for main effects of GM and sex. Two-way ANOVA followed by *Tukey’s post hoc* was used to test Chao-1 richness for main effects of GM and treatment. Since it was not possible to include a male dam group, the main effect of sex was excluded from Chao-1 analysis. Chao-1 richness was calculated using PAST 4.03 software^61^. Differences in family- and genus-level relative abundance between GMs was assessed using Wilcoxon Rank-Sum tests with a Benjamini-Hochberg correction for multiple comparisons. Due to uniform lack of normality across metabolites, a Mann-Whitney U test was used to test for differences in metabolite concentrations between GM^Low^ and GM^High^ treatment groups. Spearman’s rank correlation was used to test correlations between genus-level abundance and statistically significant metabolites. Two-way ANOVA followed by Tukey’s *post hoc* was used to test for main effects of GM and sex in the gene expression data. A student’s t-test was used to test for differences in the GM^Low^ and GM^High^ groups in the *bsh* metagenomic and metatranscriptomic read counts. A two-way ANOVA followed by Tukey’s *post hoc* was used to test for main effects of GM and sex in the BSH activity data. All univariant data analysis was performed using SigmaPlot 14.0 (Systat Software, Inc, San Jose, CA). Shapiro-Wilk test was used to test for normality using SigmaPlot 14.0. Two-way PERMANOVA testing was based on Bray-Curtis dissimilarities using the *adonis2* library from the *vegan* library^62^.

## Results

### Taxonomic differences are associated with different levels of biologically relevant metabolites

Compositional differences between GM^Low^ and GM^High^ have been described previously^36, 37, 63^, including greater richness in GM^High^ compared to GM^Low^ (**Fig. 1A**). Families enriched in GM^Low^ included *Erysipelatoclostridiaceae*, *Erysipelotrichaceae*, *Sutterellaceae*, *Saccharimonadaceae*, *Acholeplasmataceae*, and unresolved members of orders *Tissierellales* and RF39; while families enriched in GM^High^ included *Prevotellaceae*, *Marinifilaceae*, *Clostridiaceae*, *Desulfovibrionaceae*, *Deferribacteraceae*, and an unresolved family within the order *Rhodospirillales* (**Table S2**). Genera that were enriched in GM^Low^ included *Anaeroplasma, Lachnoclostridium,* and *Oscillospira*; while genera that were enriched in GM^High^ included *Odoribacter, Alloprevotella, Rikenella, Bilophila*, and *Desulfovibrio* (**Fig. S1**; **Table S3**). To determine whether these phylogenetic differences were associated with different metabolite profiles, fecal samples were collected from both male and female GM^Low^ and GM^High^ mice, and a combination of mass spectrometry (MS)-based platforms were used to measure short-chain fatty acids; branched chain fatty acids, branched chain keto-acids, and other lipids; unconjugated primary and secondary bile acids; all proteinogenic amino acids and several nonproteinogenic and acetylated amino acids; tryptophan derivatives including kynurenine, serotonin, and several indoles; B vitamins; dicarboxylic acids; glucose, fructose, and other compounds within glycolysis; all ribonucleosides and nitrogenous bases; compounds within the pentose phosphate pathway; compounds within the classic TCA cycle; and other biologically relevant microbial metabolites. Analysis of fecal metabolites revealed differences in sugar molecules involved in glycolysis, multiple amino acids, and primary bile acids (**Fig. 1B**; **Table S4**). Specifically, glucose-6-phosphate, fructose-6-phophate, ribulose-5-phosphate, and β- alanine were enriched in GM^Low^ feces, while cysteine, succinate, lactate, chenodeoxycholic acid (CDCA) and deoxycholic acid (DCA) were enriched in GM^High^ feces. These differences were particularly apparent in the feces of male mice (**Fig. S2A**), while female mice also showed a single secondary bile acid (lithocholic acid, LCA) enriched in the feces of GM^Low^ (**Fig. S2B**). When genus-level taxonomic abundances were compared to each of the significant metabolites, numerous significant correlations were identified indicating that taxonomic features in the GM were strongly associated with the differentially abundant metabolites (**Fig. S3**; **Table S5**). Metabolites were also measured in serum, collected at the same time as fecal samples. Remarkably, the only significant difference in the serum was a primary bile acid (CDCA) (**Fig. 1C**; **Table S6**). There were also several bile acids, both primary and secondary, that while not reaching statistical significance, had greater levels within the serum of female GM^Low^ and male GM^High^ mice (**Fig. S2C-D**).

**Figure 1.**
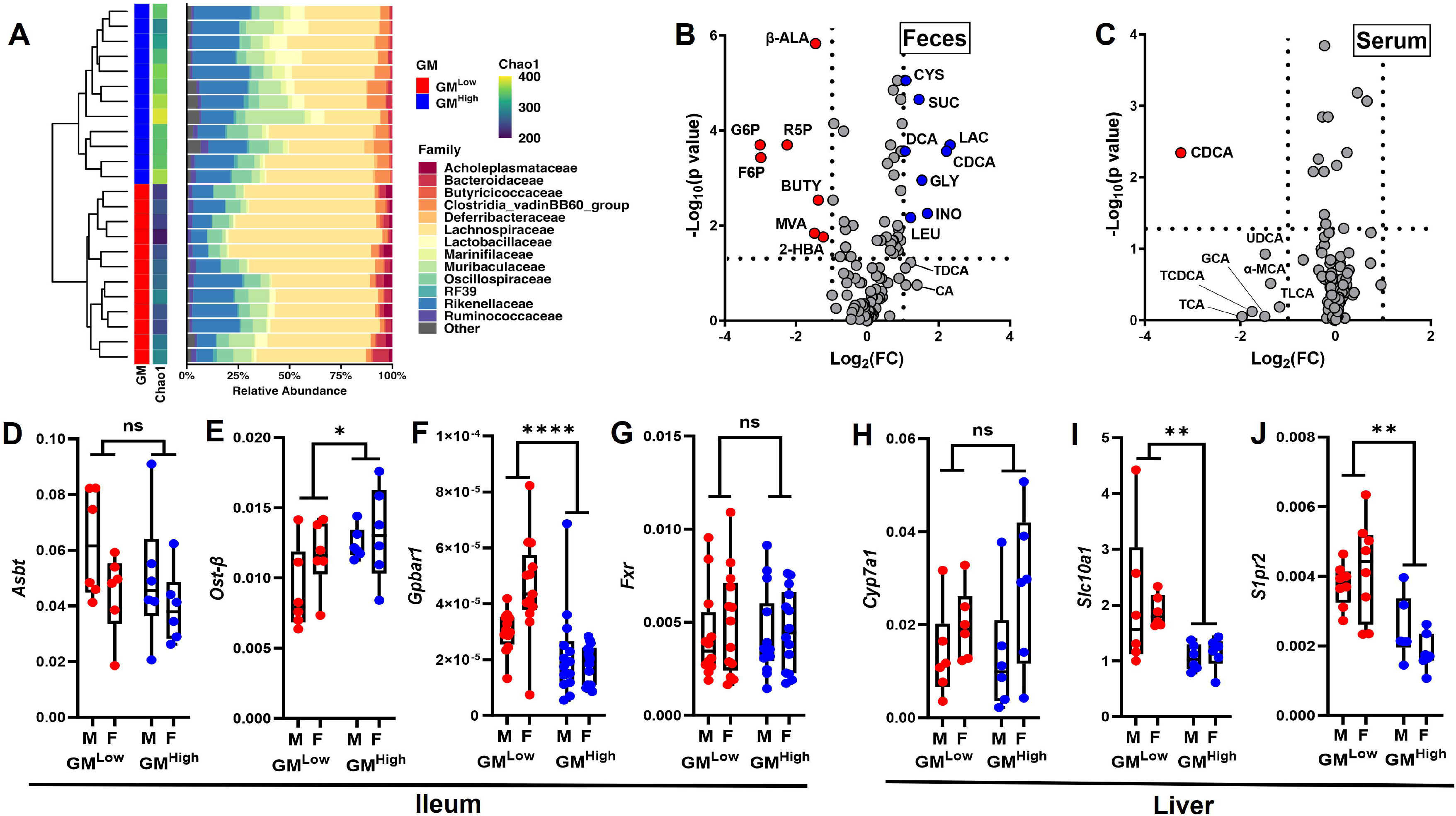
Microbiomes linked to fetal programming of complex behaviors differ in synthesis or catabolism of several metabolites including bile acids. **(A)** Hierarchical clustering of data from adult GM^Low^- or GM^High^-colonized mice, demonstrating segregation of GMs by richness and composition (legend at right). Volcano plots showing metabolites enriched in GM^Low^ (red dots) or GM^High^ (blue dots) **(B)** feces or **(C)** serum. Horizontal dashed line indicates significance of p<0.05 between GM^Low^ and GM^High^ mice by Wilcox rank sum test. Normalized ileal expression of *Asbt* **(D)**, *Ost*β **(E)**, *Gpbar1* **(F)**, and *Fxr* **(G)** male (M) and female (F) mice colonized with GM^Low^ (red dots) or GM^High^ (blue dots). Normalized hepatic expression of *Cyp7a1* **(H)**, *Slc10a1* **(I)**, and *S1pr2* **(J)** in the same groups of mice. Gene expression normalized to β*-Actin* expression. Results indicate main effect of GM in one- or two-way ANOVA, following correction for multiple tests. * p<0.05, ** p<0.01, **** p<0.0001. Abbreviations: 2-HBA (2-hydroxybutyrate), α-MCA (alpha-Muricholic Acid), β-ALA (beta-Alanine), BUTY (Butyrate), CA (Cholic Acid), CDCA (Chendeoxycholic Acid), CYS (Cysteine), DCA (Deoxycholic Acid), F6P (Fructose-6-Phosphate), G6P (Glucose-6-Phosphate), GCA (Glycocholic Acid), GLY (Glycerate), INO (Inositol), LAC (Lactate), LEU (Leucine), MVA (Mevalonic Acid), R5P (Ribulose-5-Phosphate), SUC (Succinate), TCA (Taurocholic Acid), TCDCA (Taurochenodeoxycholic Acid), TDCA (Taurodeoxycholic Acid), TLCA (Taurolithocholic Acid), UDCA (Ursodeoxycholic Acid).

To determine whether these differences in bile acids are associated with differences in bile acid cellular transport and receptor signaling, quantitative RT-PCR was performed with mRNA from both ileal and hepatic tissue. *Asbt*, a gene involved in transporting bile acid from the ileal lumen into the ileal epithelium did not show a difference in gene expression (**Fig. 1D**). However, *Ost-*β, a gene involved in transport of bile acids from the ileal epithelium into vascular circulation, showed greater expression in ileum of GM^High^ mice (**Fig. 1E**). *Gpbar1*, a G protein-coupled receptor (GPCR) also known as TGR5, was found to have greater expression in the ileum of GM^Low^ mice (**Fig. 1F**). *Fxr*, a gene critically involved in regulation of hepatic bile acid synthesis, showed no difference in ileal expression (**Fig. 1G**). Accordingly, *Cyp7a1*, a gene downstream of ileal Fxr signaling that encodes the rate-limiting protein in bile acid synthesis, also showed no difference in hepatic expression (**Fig. 1H**). Expression of *Slc10a1,* a bile acid transport protein, was higher in the liver of GM^Low^ mice (**Fig. 1I**). *S1pr2*, a GPCR activated by bile acids, was also expressed at a greater level in the liver of GM^Low^ mice (**Fig 1J**). These data indicated that the differential abundance of BAs detected in feces and serum of GM^Low^- and GM^High^-colonized mice are also associated with differential ileal and hepatic expression of bile acid receptors and transporters.

We next examined whether bile salt hydrolase (BSH), the bacterial enzyme used to deconjugate bile acids, levels showed differential expression or activity between GM^Low^- and GM^High^- colonized female mice. While no difference was detected in total gene read count (**Fig. S2E**), microbial expression of *bsh* was significantly higher in GM^Low^ (**Fig. S2F)**. The BSH family of enzymes is widely expressed among gut bacteria, and prone to significant variations in structure with some isoforms exhibiting different deconjugation activity toward various bile acids and possessing various levels of enzymatic activities^64, 65^. Using a panel of highly sensitive bioluminescent assays^48^, we compared BSH enzymatic activities of both microbiomes toward various bile acids. Our results demonstrated significantly greater BSH activity specific for cholic acid in GM^Low^ (**Fig. S2G**), but no other differences were detected in BSH activities towards other bile acids examined (**Fig. S2H-K**). Collectively, the differences in bacterial *bsh* expression and fecal and serum bile acid levels suggest greater uptake by GM^Low^ mice and greater fecal loss by GM^High^ mice. This is supported by greater hepatic expression of bile acid receptors and transporters.

### Complex microbiome-dependent behaviors determined by the parental microbiome

Prior work has revealed reproducible differences in behavior and growth between GM^Low^- and GM^High^-colonized mice^36, 37^. To determine the developmental period in which these phenotypic differences are established, we used an experimental design with four groups, comprising mice born to dams harboring GM^Low^ or GM^High^ and remaining with their birth dams until weaning (control), or cross-fostered at birth to nursing dams harboring the reciprocal microbiome (**Fig. 2A**). These groups are denoted as CF^Low^ and CF^High^, with the microbiome designation indicating the postnatal offspring composition acquired via cross-foster (CF). Comparisons were then made between the two control groups, and between the two CF groups in behavior tests associated with anxiety-related behavior and voluntary activity between five and seven weeks of age, and body weight (BW) at three and seven weeks of age.

**Figure 2.**
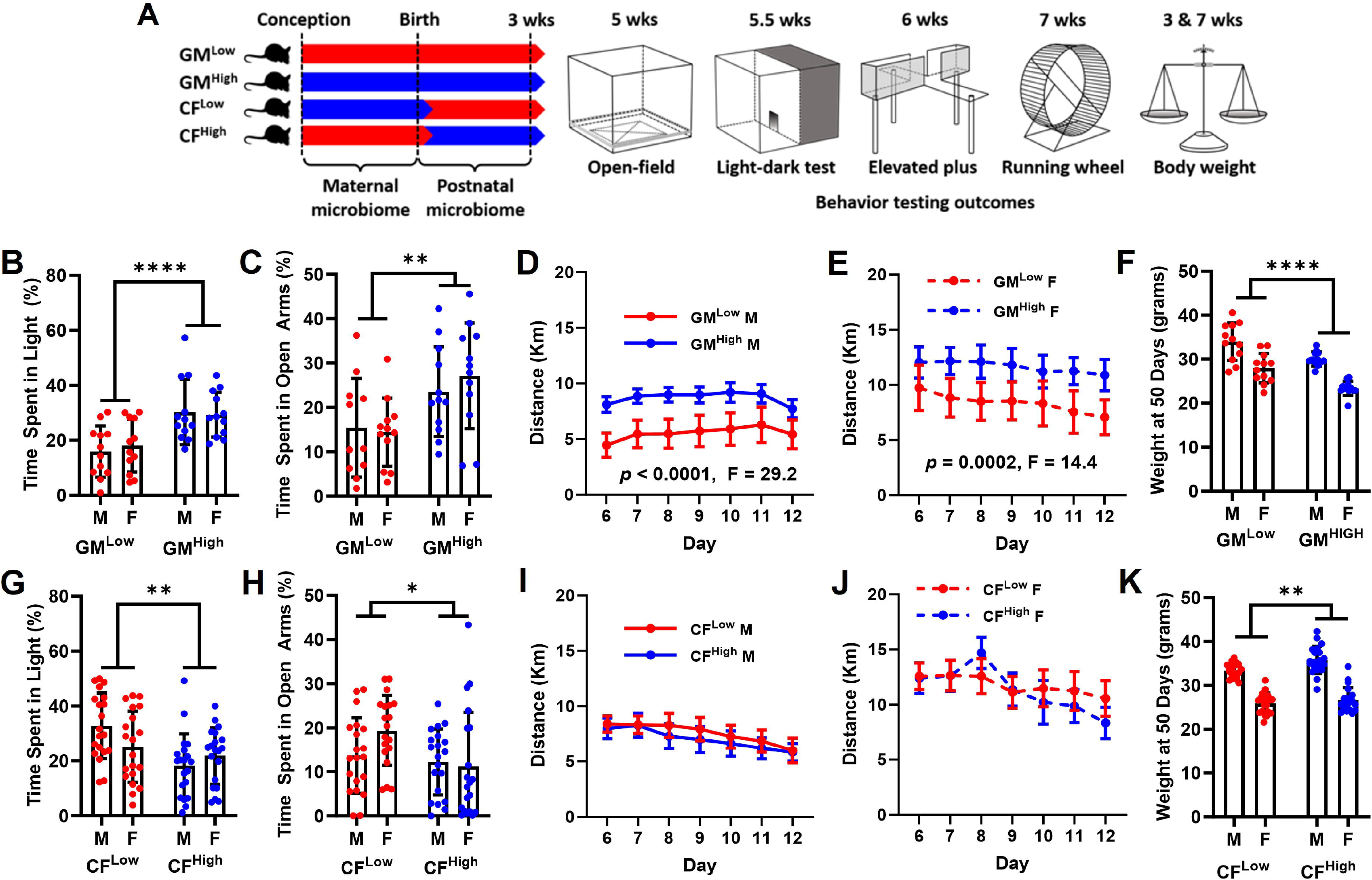
Anxiety-related behavior and other outcomes in offspring are influenced by the maternal gut microbiome. **(A)** Schematic showing the experimental groups and timing of outcome measures. Behavior outcomes in male (M) and female (F) mice colonized with GM^Low^ or GM^High^, including (**B**) time spent in the light portion of a light/dark test, (**C**) time spent in the open arms of an elevated plus maze, and distance run per day (Km) by (**D**) male and (**E**) female mice. (**F)** Body weight per mouse (grams) at day 50 in control mice. (**G-J)** Behavior outcomes in CF^Low^ and CF^High^ mice as those shown in panels **B-E**. **(K)** Body weight per mouse at day 50 in cross-fostered mice. p and F values represent effect of GM in two-way ANOVA with Tukey post hoc. Significant sex-dependent differences were detected in distance on running wheels and body weight in both control and cross-fostered mice, with no significant interactions between GM and sex in any tests. * p<0.05, ** p<0.01, **** p<0.0001

In agreement with previous studies^37^, age-, sex-, and genotype-matched CD-1 mice colonized with GM^Low^ spent less time and traveled less distance in the light portion of a light-dark transition (LDT) test (**Fig. 2B**, **Fig. S4A**), and the open arms of an elevated plus maze (EPM) test (**Fig. 2C**, **Fig. S4B**), relative to mice colonized with GM^High^, indicating differential effects on anxiety-related behavior of the two microbiomes. No behavioral differences were observed in the open-field exploration test between GM^Low^ and GM^High^ mice (**Fig. S4C-D**). To assess voluntary physical activity, mice were singly housed with bluetooth wireless running wheels for a five-day acclimation period followed by a seven-day test period. Both male (**Fig. 2D**) and female (**Fig. 2E**) mice colonized with GM^High^ ran significantly more than mice with GM^Low^ (*p* < 0.0001, F = 29.2, and *p* = 0.0002, F = 14.4, respectively). Previously reported differences in body weight (BW) at weaning and adulthood were also reproducible in the GM^Low^ and GM^High^ groups^37^ (**Fig. 2F**, **Fig. S4E**). Collectively, these data confirmed microbiome-associated differences in anxiety-related behavior, voluntary physical activity, and body weight, in sex-, age-, and genotype-matched mice.

Microbial 16S rRNA amplicon sequencing was used to confirm that CF mice harbored a microbiome of comparable richness and beta-diversity to that present in surrogate dams, in both directions of CF (**Fig. S5A-C**). Behavioral analysis showed the GM-associated differences in the LDT, independent of sex, were reversed in the comparison between CF^Low^ and CF^High^ mice, with the CF^High^ mice demonstrating behavior suggestive of greater anxiety (**Fig. 2G**, **Fig. S4F**). Similarly, the robust GM-dependent differences in the EPM were reversed in the CF^Low^ and CF^High^ mice (**Fig. 2H**, **Fig. S4G**) indicating that the birth dam microbiome has a substantial, if not dominant, influence on neurodevelopmental events contributing to anxiety-related behavior in the offspring. While a significant difference of total distance traveled in the OFE test was observed, distance traveled in the center was not found to be significant in the cross-foster groups **(Fig. S4H-I)**. The significant GM-associated differences in voluntary physical activity were absent in male and female CF mice (**Fig. 2I-J**). Comparison of BW revealed no difference at weaning (**Fig. S4J**) and reversal in adulthood (**Fig. 4K**), such that CF^High^ mice weighed more than age-matched CF^Low^ mice. Collectively, these data supported an equivalent or dominant effect of the birth dam GM on offspring behavioral phenotypes and body weight.

**Figure 3.**
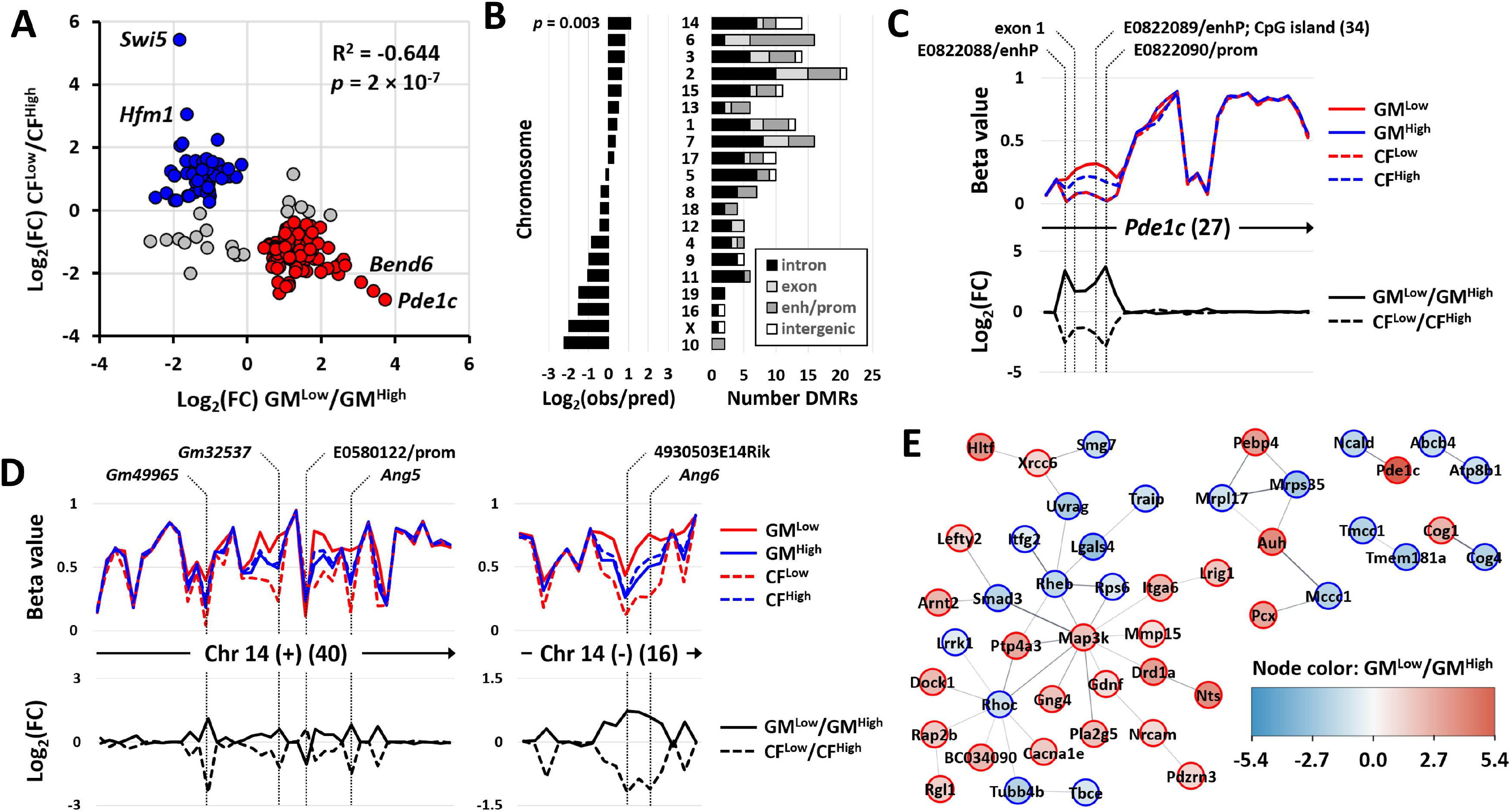
Maternal microbiome is associated with gene methylation in offspring hippocampus. **(A)** Dot plot showing normalized difference [Log_2_(FC)] in mean methylation between GM^Low^ and GM^High^, and between CF^Low^ and CF^High^, of all CpG markers achieving a Log_2_(FC) > 1 in either comparison. **(B)** Ratio of observed to predicted differentially methylated regions (DMRs), and locations of DMRs relative to gene bodies, on each chromosome. **(C)** Mean beta values of 27 CpG sites spanning the enhancers, promoters, introns and exons across the *Pde1c* gene in all four groups, with specific sites indicated above (upper); and log_2_(FC) between control and cross-fostered groups (lower). Arrows on X-axis indicate direction of transcription. Numbers on the X-axis indicated number of CpG sites analyzed. **(D)** Mean beta values (upper) and Log_2_(FC) between groups (lower) across adjacent regions of the positive (left) and negative (right) strand of a region of chromosome 14 containing genes for *Ang5*, *Ang6*, and other genes. **(E)** Protein interaction networks among products of DMRs, with node color indicating Log_2_(FC) in methylation between GM^Low^ and GM^High^.

**Figure 4.**
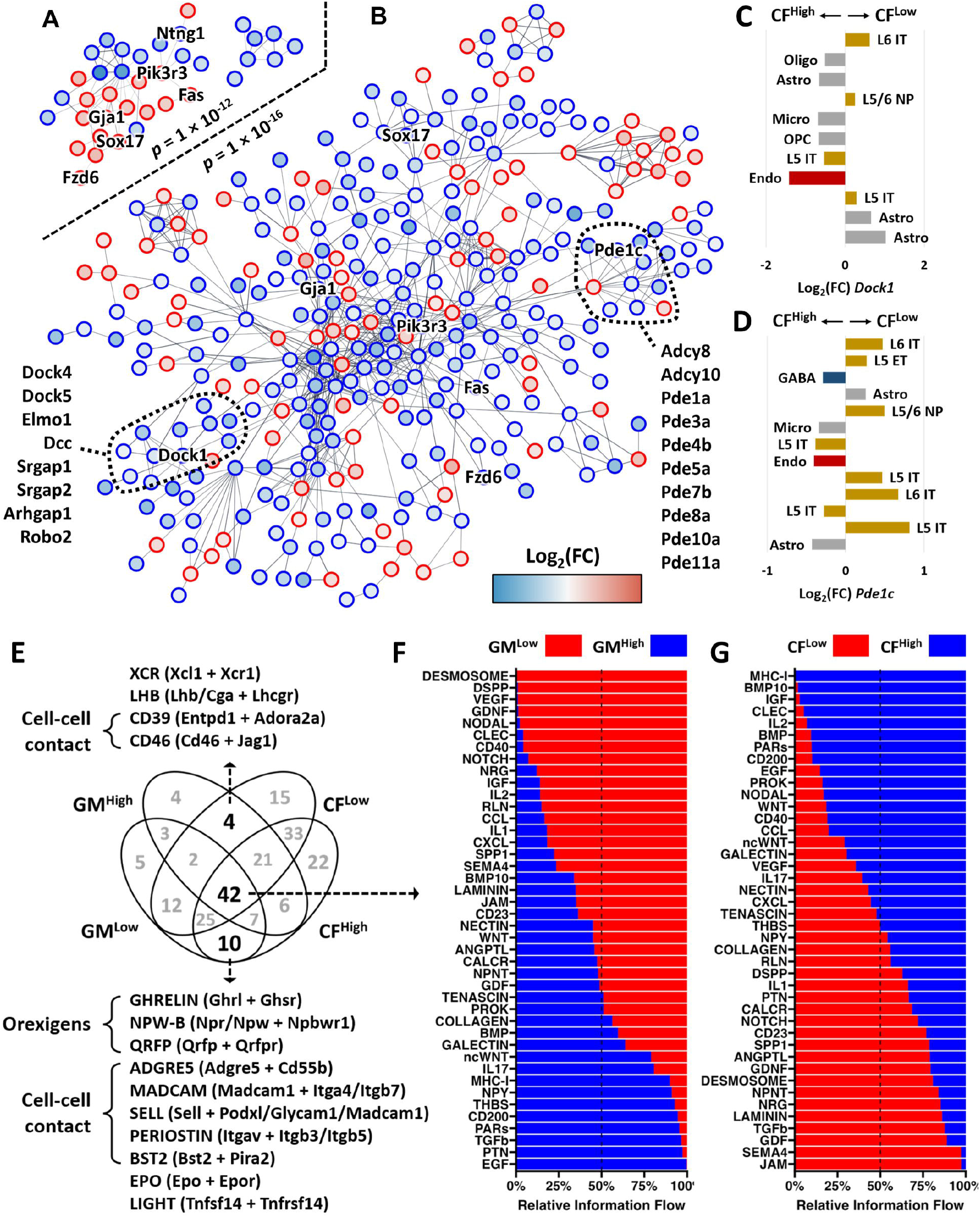
Interaction networks constructed using differentially expressed genes (DEGs) identified in hippocampal endothelial and perivascular cells of control. (**A**) and cross-fostered (**B**) mice. Highlighted nodes include DEGs identified in both comparisons and showing a pattern of fetal imprinting, and genes that are also differentially methylated (*Pde1c*, *Dock1*), or closely related (PDE cluster, *Dcc*, *Elmo1*, *Dock4*, *Dock5*). **c, d.** Log_2_FC in expression of (**C**) Pde1c and (**D**) Dock1 in hippocampal cell clusters of male and female CF^Low^ and CF^High^ mice, as determined via real-time snRNAseq. **(E)** Venn diagram showing number of inferred cell-to-cell communication pathways identified in hippocampus of GM^Low^, GM^High^, CF^Low^, and CF^High^ mice. Pathways listed above and below the diagram were selectively identified in the indicated groups. Bar charts showing the relative degree of cell-to-cell communication between (**F**) GM^Low^ and GM^High^, and (**G**) CF^Low^, and CF^High^ mice in the indicated pathways.

### Fetal programming of gene methylation in hippocampus by the parental microbiome

Reasoning that an influence on offspring behavior by the parental microbiome must have a biological foundation in the CNS, and a mechanism by which cellular function and gene expression are established during fetal development, we next examined gene expression and its epigenetic regulation in offspring hippocampus, given its central role in anxiety-related behavior^66–71^. To identify effects of the parental microbiome on fetal programming of DNA methylation in the hippocampus, we performed genome-wide DNA methylation profiling in female mice. Analysis of methylation across the entire array, and parallel comparisons between samples from GM^Low^ or GM^High^ mice, and from CF^Low^ or CF^High^ mice, identified only 196 differentially methylated regions (DMRs) with beta values differing by log_2_FC > 1 (**Table S7**). Remarkably, 176 of those 196 sites (89.8%) showed a difference in methylation at the same site in the reciprocal contrast, such that offspring methylation matched that of their birth dam (**Fig. 3A**, **R^2^ = -0.644, *p* = 2 × 10^-7^**). DMRs identified in offspring hippocampi were distributed across the genome, occurring most often early in gene bodies or enhancer/promoter regions and roughly correlating in frequency to chromosomal gene content, with the exception of an apparent enrichment for DMRs on chromosome 14 (**Fig. 3B, *p* = 0.003**). Differential methylation was detected at five contiguous markers mapped to promoters, enhancers, and exon 1 of the *Pde1c* gene, encoding phosphodiesterase 1c, a regulator of Ca^2+^ and cGMP-dependent intracellular signaling. This methylation pattern differed between hippocampus from GM^Low^ and GM^High^ mice, and was conserved between birth dam and offspring, regardless of postnatal colonization (**Fig. 3C**). Similarly, microbiome-associated differences in methylation were identified at multiple closely spaced markers on both strands of chromosome 14, including markers associated with the genes *Ang5* and *Ang6* (**Fig. 3D**), encoding members 5 and 6, respectively, of the angiogenin, ribonuclease A family.

To identify shared pathways or commonalities among the functional products (i.e., proteins) encoded by protein-coding genes among the DMRs, a STRING analysis was performed using a final input of 144 gene names^72^. Interaction analysis resulted in assembly of one large network with 32 nodes, a smaller network with six nodes, and four dyads (**Fig. 3E**). While enrichment analysis failed to detect greater network connectivity than would occur at random, stratified analysis of the 32 nodes in the large network revealed 10 significantly enriched Gene Ontology (GO) Biological Processes; five GO Molecular Functions including TGF-β receptor binding (GO:0005160, strength 1.94, FDR-adjusted *p* = 0.015) and GTPase activity (GO:0003924, strength = 1.09, FDR-adjusted *p* = 0.042); five KEGG pathways including Gap junction (mmu04540, strength = 1.38, FDR-adjusted *p* = 0.033) and TGF-β signaling pathway (mmu04350, strength = 1.32, FDR-adjusted *p* = 0.033); and four Reactome pathways including axon guidance (MMU-422475, strength = 1.08, FDR-adjusted *p* = 0.048). Network STRING analysis results provided in **Table S8**.

### Fetal programming of gene expression in hippocampus by parental microbiome

We next performed single nuclei RNAseq on hippocampus collected from control mice and cross-fostered mice. Based on expression of cell-specific markers, 11 cell clusters were identified in the control mice (**Fig. S6A-B**) and 24 cell clusters were identified in the cross-fostered mice (**Fig. S6C-D**). Gene expression in GM^Low^ and GM^High^ control mice was compared using generalized additive regression models, and differentially expressed genes (DEGs) within each treatment group were identified (**Table S9**). Similarly, mice from CF^Low^ and CF^High^ were examined for DEGs within each cell cluster (**Table S10**). After DEGs from each cell cluster had been determined using a cutoff magnitude difference of Log_2_FC > 1.5, STRING analysis was used to determine the protein-protein interactions of the DEGs from each treatment^72^. The number of DEGs from each cell cluster were compared to the mean node degree received from the STRING analysis results to determine which cell clusters contained a high number of DEGs that were most likely to interact within protein pathways. Interestingly the cell cluster with the highest number of DEGs, as well as the highest number of mean node degree of protein-protein interactions, in both control and cross-foster mice was identified as hippocampal endothelial cells (**Fig. S7A-B**). STRING analysis was used to generate interaction networks using DEGs identified in the endothelial cells of control mice as well as the cross-fostered mice (**Fig. 4A-B**). Of these interactions, six DEGs (*Fas, Fzd6, Gja1, Ntng1, Pik3r3, Sox17*) showed a pattern of fetal programming. Of note, numerous DEGs (including *Pde1c*, *Dock1, and Pdzrn3*) identified were also found to be differentially methylated or closely related to differentially methylated genes. When L5 IT glutamatergic neurons and astrocytes were examined using interaction networks, they also contained multiple DEGs that showed a pattern of fetal programming as well as DEGs that were identified as differentially methylated (**Fig. S8A-B**; **Fig. S9A-B**). When we examined the Log_2_FC of *Dock1* expression of CF^Low^ and CF^High^, *Dock1* was shown to be upregulated in the clusters of endothelial cells, oligodendrocyte, microglial cells, and subsets of astrocytes and glutamatergic neurons of CF^High^ mice compared with CF^Low^ (**Fig. 4C**). Similarly, when we examined expression of *Pde1c*, it too was found to be increased in the endothelial cell cluster of CF^High^ mice, though not consistent with the expression patterns of other cells seen with *Dock1* (**Fig. 4D**). We next used CellChat software^59^ to impute cell-cell communication via patterns of coordinated ligand-receptor expression in each control and cross-foster group, to identify cell signaling pathways that show a pattern of fetal programming. There were 42 cell-cell communication pathways that were shared among the control mice and the cross-foster mice (**Fig. 4E**). Of the 42 pathways identified, 15 showed a pattern of fetal programming including VEGF, IGF, IL2, TGFβ, WNT, and NPY (**Fig. 4F-G**). Interestingly, three cell-cell communication pathways that were identified in only GM^Low^ and CF^High^ were appetite-stimulating orexigenic neuropeptide pathways.

## Discussion

Studies comparing germ-free and SPF mice demonstrate that the parental microbiome can affect offspring phenotypes associated with neurodevelopment^73^, metabolic diseases including obesity^74^, and organogenesis in the CNS and intestines^75^. While challenging to study in human cohorts, recent retrospective analyses suggest a dominant influence of the maternal microbiome on offspring phenotypes related to asthma^76^, neurodevelopment^77^, and metabolic diseases including obesity and diabetes^78^. The current data demonstrate that differences among native parental microbiomes can influence neurodevelopment and behavioral outcomes in the offspring. The current findings and prior studies^36, 37^ show reproducible effects of these native SPF microbiomes on certain phenotypes. GM^Low^-colonized CD-1 mice are consistently heavier than age- and sex-matched mice colonized with GM^High^, and the same effect is observed in inbred C57BL/6J and BTBR T^+^ Itpr3^tf^/J mice^79^. Behavior and BW data from the cross-foster (CF) mice provide strong evidence of a dominant effect of the parental GM on these behavioral phenotypes in offspring. While we did not measure food intake in the current study, previous work showed that the heavier GM^Low^-colonized mice consume more food (normalized to BW) than age- and sex-matched GM^High^-colonized at all timepoints examined^36^. The differences in BW between CF and control offspring would suggest that these feeding behaviors are similarly programmed during fetal, embryonic, or even pre-fertilization events. As such, these findings raise the possibility of a connection between the anxiety-related behaviors and the behaviors underlying the difference in BW and voluntary physical activity, representing a constellation of behavioral phenotypes influenced by common features within the parental microbiomes.

The GM can influence host physiology through microbially derived metabolites in peripheral circulation^73^, interactions with the immune system^80^, and stimulation of the vagus nerve or enteric nervous system by microbially derived neurotransmitters^81^ and other molecules. Gut metabolites have been implicated as a means by which the parental microbiome can influence fetal development^73, 82^. The present data provide evidence of a functional difference between these native SPF microbiomes, including differential abundance of several bile acids. Bile acids stored in the gallbladder are released into the duodenum following food intake, and the observed differences in fecal bile acids may reflect differences in *bsh* expression, food intake, or other factors. Regardless, the observed differences in ileal and hepatic bile acid transporters and receptors indicate that the differences in bile acid levels are physiologically relevant to the host. The observed difference in expression of *Slc10a1 (Ntcp)* may reflect a mechanism to regulate reabsorption of bile acids. GM^Low^ mice also demonstrated greater hepatic expression of *S1pr2*, a GPCR that when bound to primary conjugated bile acids is involved in the regulation of hepatic lipid metabolism^83^. There is considerable interest in the role of bile acids in anxiety and depressive disorders^84, 85^, and causative links have been shown between bile acids, bile acid receptor signaling, and these outcomes^86–89^. CDCA, present at greater levels in the serum of GM^Low^-colonized females, has been shown to readily cross the blood brain barrier^90^ and influence the expression of transcription factors CREB and BDNF through FXR activation, which when down-regulated can lead to decreased neuroplasticity and mood disorders including anxiety^91^. The current findings suggest that differences in the native microbiome, independent of dietary challenge or host insult, can have intergenerational effects on these outcomes.

Analyses of hippocampal DNA methylation and gene expression were performed to document and compare the effects of the parental or offspring GM on those processes, and identify specific genes, pathways, and cell types associated with the observed phenotypic differences. It is well-established that the GM can influence the epigenome of the host^92–94^. The distribution and relationship of genes affected by differential methylation reflects a semi-stochastic effect across the genome, with enrichment of genes and pathways associated with TGF-β signaling and GTPase activity, both of which were identified in the single-cell transcriptome data as well. Four of the 144 protein-coding genes (*Dlx5*, *Drd1*, *Zfp64*, and *BC034090*) identified as DMRs here are included among a comprehensive list of 384 genes known to undergo fetal imprinting^95^. This suggests that the GM-associated effects on methylation of those and perhaps other DMRs occurred in germline cells pre-fertilization. Indeed, several recent studies have revealed the influence of the paternal microbiome on germline methylation and offspring outcomes^96–98^. As all matings in the current study were between mice sharing the same microbiome, it is unclear whether the effects of the GM on offspring DNA methylation occurred pre- or post-fertilization and whether the maternal lor paternal microbiome had a dominant or selective influence. As even transient co-housing to breed mice results in sharing of the GM, *in vitro* fertilization or similar methods would be needed to investigate those questions. There was an incredibly high degree of conservation across all mice in the degree of methylation at the vast majority of CpG sites included in the BeadChip array. However, we also observed a high degree of conservation between dams and offspring in the specific DMRs affected by the GM, suggesting the affected loci are not the result of random DNA methyltransferase (DNMT) activity, but rather an outcome with a teleological explanation. While speculative, the methylation and gene expression profiles following a pattern of fetal programming may represent an intergenerational feedback mechanism wherein nutrient availability in the parent may program the trafficking of, or receptor response to, microbial metabolites as a way of fine-tuning offspring metabolism.

It is worth noting that the number and connectivity of DEGs were greatest in endothelial cells, in both control and CF mice. These cells supply blood to tissues within the CNS and comprise the blood-brain barrier. The greater size of GM^Low^-colonized mice would necessitate greater amount of peripheral vasculature to adequately perfuse tissues. Indeed, prior work found the total cardiac weight of GM^Low^-colonized mice to be significantly greater than age- and sex-matched GM^High^-colonized mice, and no difference in cardiac weight when normalized to total BW^37^, indicating differential growth of the circulatory system commensurate with the difference in BW. Moreover, the same study found no difference in body composition based on DEXA scanning, and a significant correlation between BW and crown-to-rump length, further indicating that the observed phenotypic difference is associated with somatic growth rather than adiposity. Several of the pathways identified in the CellChat analysis showing patterns of fetal programming represented growth factors including TGF-β, vascular endothelial growth factor (VEGF) and insulin-like growth factor (IGF).

Two of the DEGs identified in the endothelial cells included *Pde1c* and *Dock1*, two genes that were also found to be differentially methylated. PDE1C is a member of the phosphodiesterase family of enzymes involved in the production of cyclic guanosine monophosphate (cGMP) and cyclic adenosine monophosphate (cAMP). The production of cAMP is necessary to maintain the integrity of the blood-brain barrier (BBB)^99^, and excessive levels of cGMP are associated with anxiety and depression^100^. DOCK1 is a protein belonging to the dedicator of cytokinesis (DOCK) family of guanine exchange factors (GEFs) involved in activation of G proteins. DOCK1 is involved in neuronal development and angiogenesis^101^. The difference in expression of *Pde1c* and *Dock1* in the endothelial cells may indicate a difference in permeability of the BBB within the hippocampus. It is also worth noting that a number of other phosphodiesterase and DOCK genes were identified as DEGs in endothelial cells, with a largely uniform direction of difference. This suggests a consistent differential effect of these microbiomes on two major mechanisms of regulating intracellular signal transduction (i.e., cyclic nucleotide generation and GEF activity) across a sizeable range of surface receptors, including GPCRs. GPCRs are among the largest classes of receptors and common drug targets^102^, responding to neurotransmitters, hormones, and a wide range of sensory cues. With widespread and strong expression in the CNS^103^, GPCRs are also broadly expressed by enteroendocrine cells^104^, vagal efferents^105^, and other cells throughout the gut^106^.

Another gene of interest found to be differentially expressed in astrocytes is the gene encoding for brain-derived neurotrophic factor (BDNF). BDNF over-expression leads to a decrease in anxiety-related behavior in mice^107^. However, we observed relatively greater *Bdnf* expression in GM^Low^ and CF^High^ mice which were found to have relatively increased anxiety-related behavior, and these findings therefore need to be explored further.

Interestingly, CellChat analysis detected coordinated expression of genes involved in orexigenic pathways including *Ghrelin*, *Npr*, and *Qrfp* only in GM^Low^ and CF^High^ mice, the two groups with greater BW. These differences in intercellular signaling may provide an explanation for the observed differences in feed intake. These findings highlight cell type specific differences in hippocampal gene expression in genes identified within DMRs and genes in pathways associated with growth and feeding behavior, giving a possible reason for the increased weight noted in these two groups.

We recognize a number of limitations within this study. For example, methylome analysis was restricted to samples from female offspring and their dams due to resource constraints. Similarly, methylome and transcriptome analyses were limited to hippocampal tissue. While these data provide strong proof-of-principle and demonstrate the utility of the experimental model, additional work is needed to determine whether the observed differences in hippocampal methylation and gene expression are conserved in other regions of the CNS or even other tissues. Lastly, the parental generation was represented by the dams in all analyses presented here. There has been growing evidence for paternal programming of offspring through the epigenome^96, 98^, and additional studies are needed to determine whether the observed effects are due to pre- or post-fertilization events..

## Conclusion

In total, the findings presented here demonstrate that features within healthy native GMs exert an intergenerational effect on offspring behavior, growth, DNA methylation, and gene expression within the central nervous system, and strongly suggest a relationship between these factors during fetal development. Moreover, these findings implicate bile acids as potential mediators of these effects, including changes in GPCR signal transduction and pathways involved in feeding behavior.

## List of abbreviations

AD –: Anxiety disorders
BSH –: Bile salt hydrolase enzyme
*bsh* –: Bile salt hydrolase gene
CF –: Cross-fostered offspring
CNS –: Central Nervous System
DEGs –: Differentially expressed genes
DMRs –: Differentially methylated regions
EPM –: Elevated plus maze test
GM –: Gut microbiome
LDT –: Light/dark transition test
OFE –: Open-field exploration test
OO –: Obesity/Overweight
snRNA-seq –: Single nuclei RNA sequencing

## Declarations

### Ethics Approval and consent to participate

All activities described here were performed in accordance with the guidelines put forth in the Guide for the Care and Use of Laboratory Animals and were approved by the Institutional Animal Care and Use Committee (IACUC) of the University of Missouri, an AAALAC accredited institution.

### Consent for publication

Not applicable

### Availability of data and material

All data supporting our analyses and reported conclusions have been deposited in the appropriate data repositories and are publicly available. Metagenomic, metatranscriptomic, 16S rRNA amplicon sequencing, and single nuclei RNAseq data are available at the National Center for Biotechnology and Informatics (NCBI) Sequence Read Archive (SRA) under the BioProject accession number PRJNA885816. Mouse Methylation BeadChip array data are available at the Gene Expression Omnibus (GEO) under accession GSE239371.

## Author contributions

K.L.G and A.C.E conceived the project and wrote the manuscript. K.L.G., Z.L.M., R.E.M., P.K., E.A.G. performed experiments and analyzed data. Z.L.M. assisted in microbiome bioinformatic analysis, and figure generation and editing. S.B.B., Z.L.M, M.R, and L.M.C performed the snRNAseq bioinformatic analysis. R.C. assisted in designing and collecting data for the methylome analysis experiment, and performed the methylome bioinformatic analysis. N.J.B. assisted in designing and collecting data for the snRNAseq experiment. P.W. contributed resources and assisted with snRNAseq analysis. D.J.D. assisted in design and performance of qRT-PCR assays. C.L.F. and J.A.L. were responsible for generating and maintaining the GM^Low^ and GM^High^ colonies of CD-1 mice used in the experiments. K.L.G., S.B.B., Z.L.M., R.E.M., P.K., N.J.B., D.J.D., M.R., L.M.C., E.A.G., J.A.L., C.L.F., P.W., R.C., and A.C.E. all assisted in editing the manuscript.

## Acknowledgements

The authors would like to thank the University of Iowa metabolomics core for assisting with metabolomic data acquisition and analysis. The authors would also like to thank the University of Missouri Mutant Mouse Resource and Research Center for donating the CD-1 mice used in this study. The authors would also like to thank Rebecca Dorfmeyer for prepping fecal DNA samples for 16S rRNA amplicon analysis.

---

**Figure S1.**
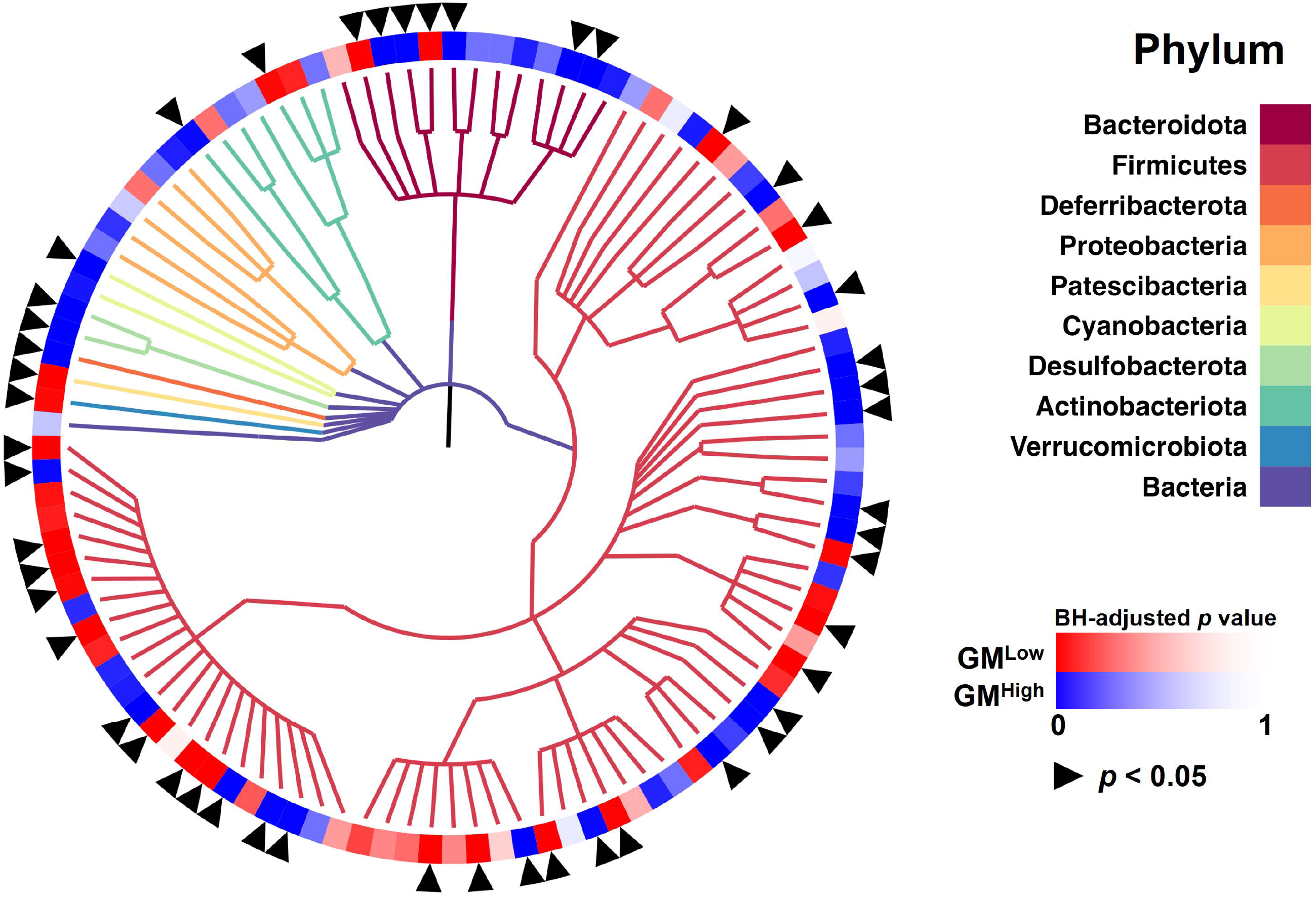
Cladogram representing genera taxa identified within each phyla. Arrows indicate genus with statistically significant abundance differences. Wilcox rank sum test with Benjamini-Hochberg corrected p values. List of genera identified provided in **Table S3**.

**Figure S2.**
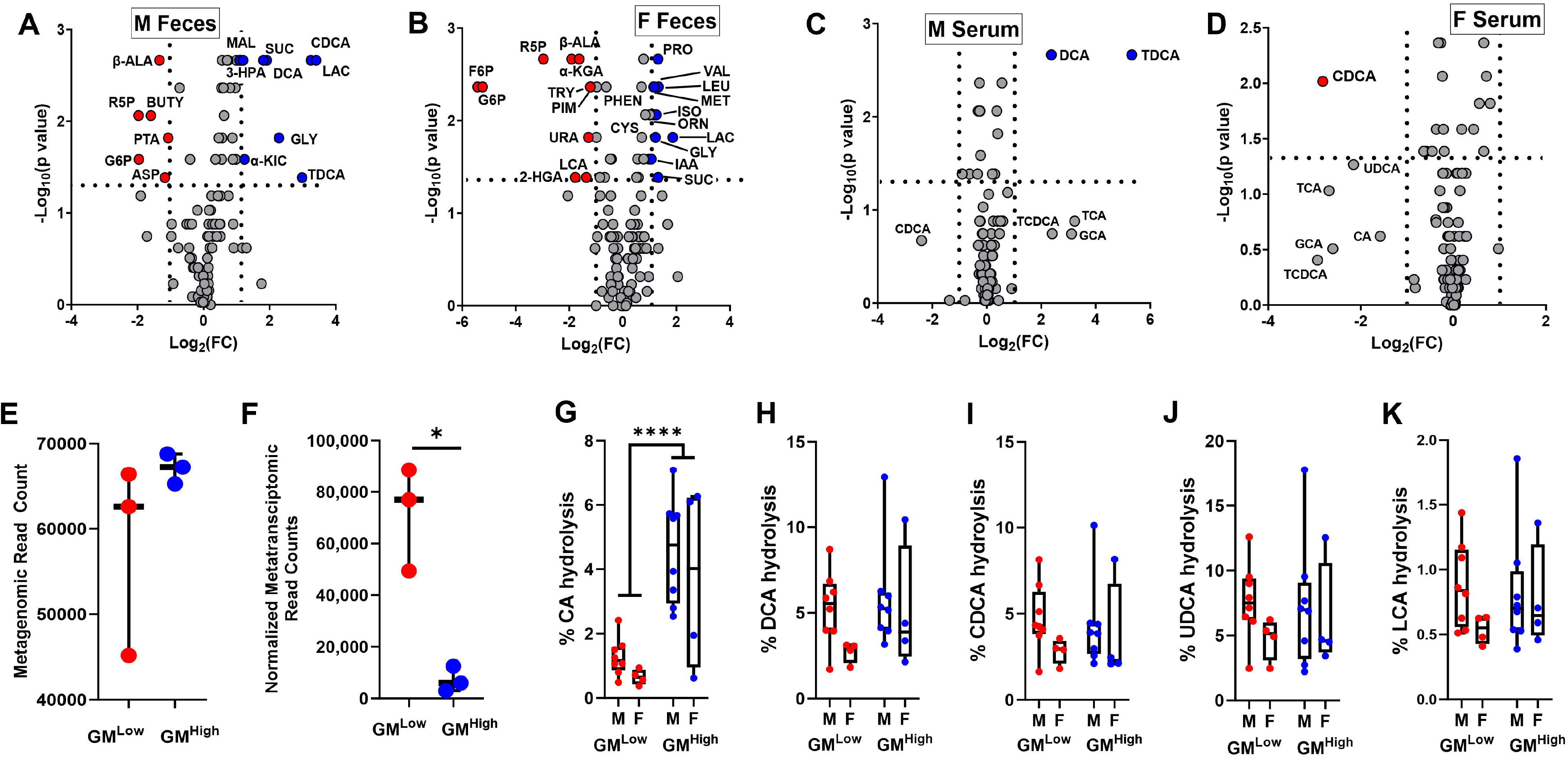
Volcano plots showing the metabolites enriched in feces of GM^Low^(red dots) and GM^High^ (blue dots) **(A)** males, and **(B)** females. Volcano plots showing the metabolites enriched in serum of GM^Low^(red dots) and GM^High^ (blue dots) **(C)** males, and **(D)** females and Dot plots representing **(E)** gene and **(F)** the normalized (transcript/gene) expression in the feces of adult female mice colonized with GM^Low^ or GM^High^ (*n* = 3/GM). Box plots and individual data showing hydrolytic activity of bile salt hydrolase (BSH) in adult male and female mice colonized with GM^Low^ or GM^High^, determined using five different bile acid-conjugated bioluminescent probes specific for **(G)** cholic acid (CA), **(H)** deoxycholic acid (DCA), **(I)** chenodeoxycholic acid (CDCA), **(J)** ursodeoxycholic acid (UDCA), and (**K**) lithocholic acid (LCA). *p<0.05, **** p<0.0001. Abbreviations: 2-HGA (2-hydroxygluterate), 3-HPA (3-Hydroxyproprionate), α-KGA (alpha- Ketogluterate), α-KIC (alpha-Ketoisocaproate), ASP (Asparagine), β-ALA (beta-Alanine), CDCA (Chendeoxycholic Acid), CYS (Cysteine), DCA (Deoxycholic Acid), F6P (Fructose-6-Phosphate), G6P (Glucose-6-Phosphate), GLY (Glycerate), IAA (Indoleacetic Acid), ISO (Isocitrate), LAC (Lactate), LCA (Lithocholic Acid), LEU (Leucine), MAL (Malate), MET (Methionine), ORN (Ornithine), PHEN (Phenylalanine), PRO (Proline), PTA (Pantothenic Acid), R5P (Ribulose-5- Phosphate), SUC (Succinate), TDCA (Taurodeoxycholic Acid), TRY (Tryptamine), VAL (Valine).

**Figure S3.**
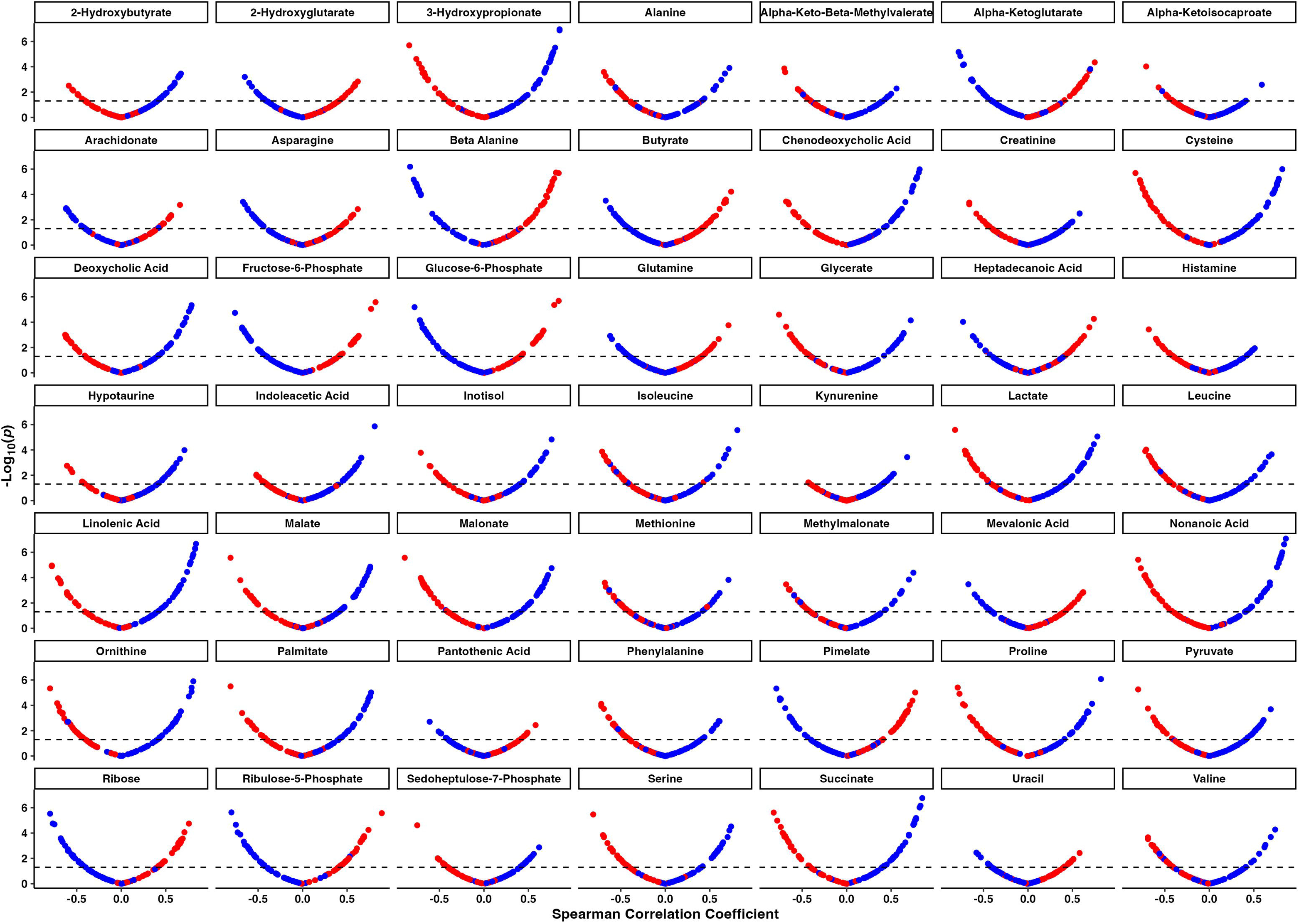
Spearman correlation coefficient plots comparing relative abundance at the genera level and metabolite concentrations to the p value of the Spearman correlation. Spearman correlation coefficient is plotted along the x-axis for each metabolite, and -Log_10_(p value) of the correlation between relative taxa abundance and metabolite concentration is plotted on the y-axis. Red dots represent genera with an increased relative abundance in GM^Low^, and blue dots represent genera with an increased relative abundance in GM^High^. Dotted lines in each plot represent statistical significance of p<0.05.

**Figure S4.**
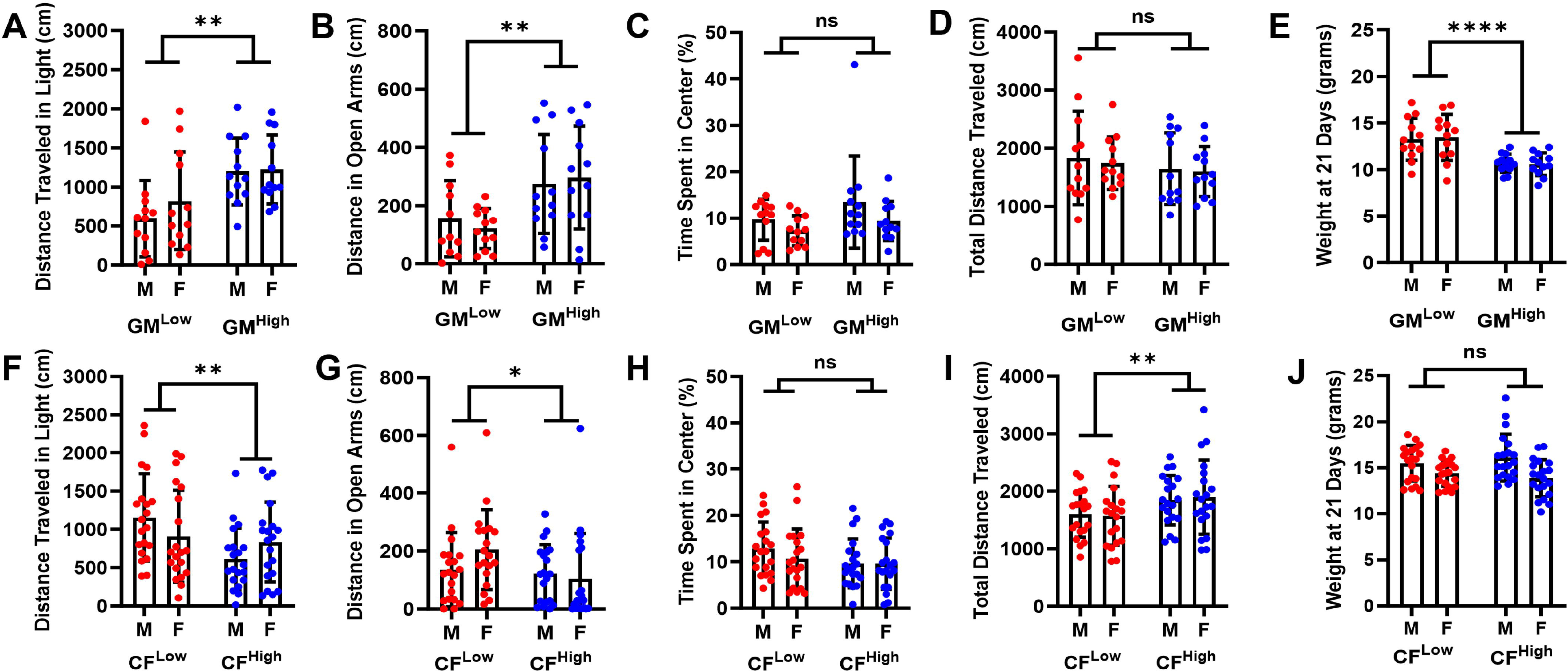
Dot and bar plots showing other results of behavior tests in adult male (M) and female (F) GM^Low^ and GM^High^ mice including (**A**) distance traveled in the light portion of the light/dark test and (**B**) open arms of the elevated plus maze. (**C**) Total distance traveled and (**D**) time spent in the center zone during OFE. (**E**) Body weight at 21 days of age of GM^Low^ and GM^High^ mice. (**F-J**) Same outcomes in CF^Low^ and CF^High^ mice as those shown in panels **A-D. (J)** Body weight at 21 days of age of CFLow and CFHigh mice. * p<0.05, ** p<0.01, **** p<0.0001.

**Figure S5.**
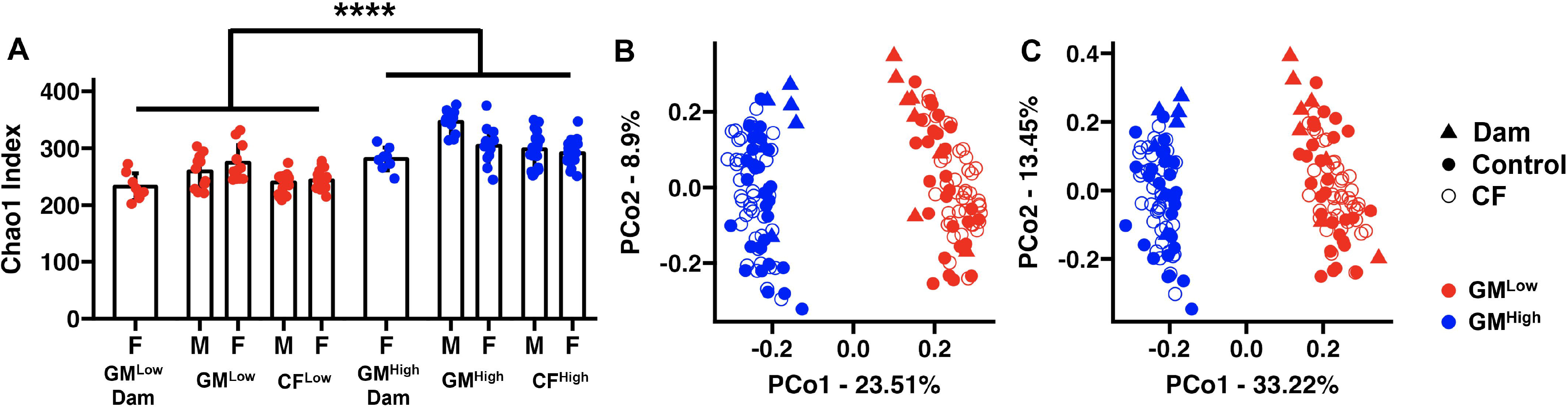
Dot and bar plots showing differences in (**A**) richness and principal coordinate analysis (PCoA) plots showing differences in beta-diversity based on (**B**) Jaccard and (**C**) Bray-Curtis distances, between adult male (M) and female (F) mice colonized with GM^Low^ or GM^High^, and similarities in all of the above metrics between CF mice at seven weeks of age and their cognate birth dams.

**Figure S6.**
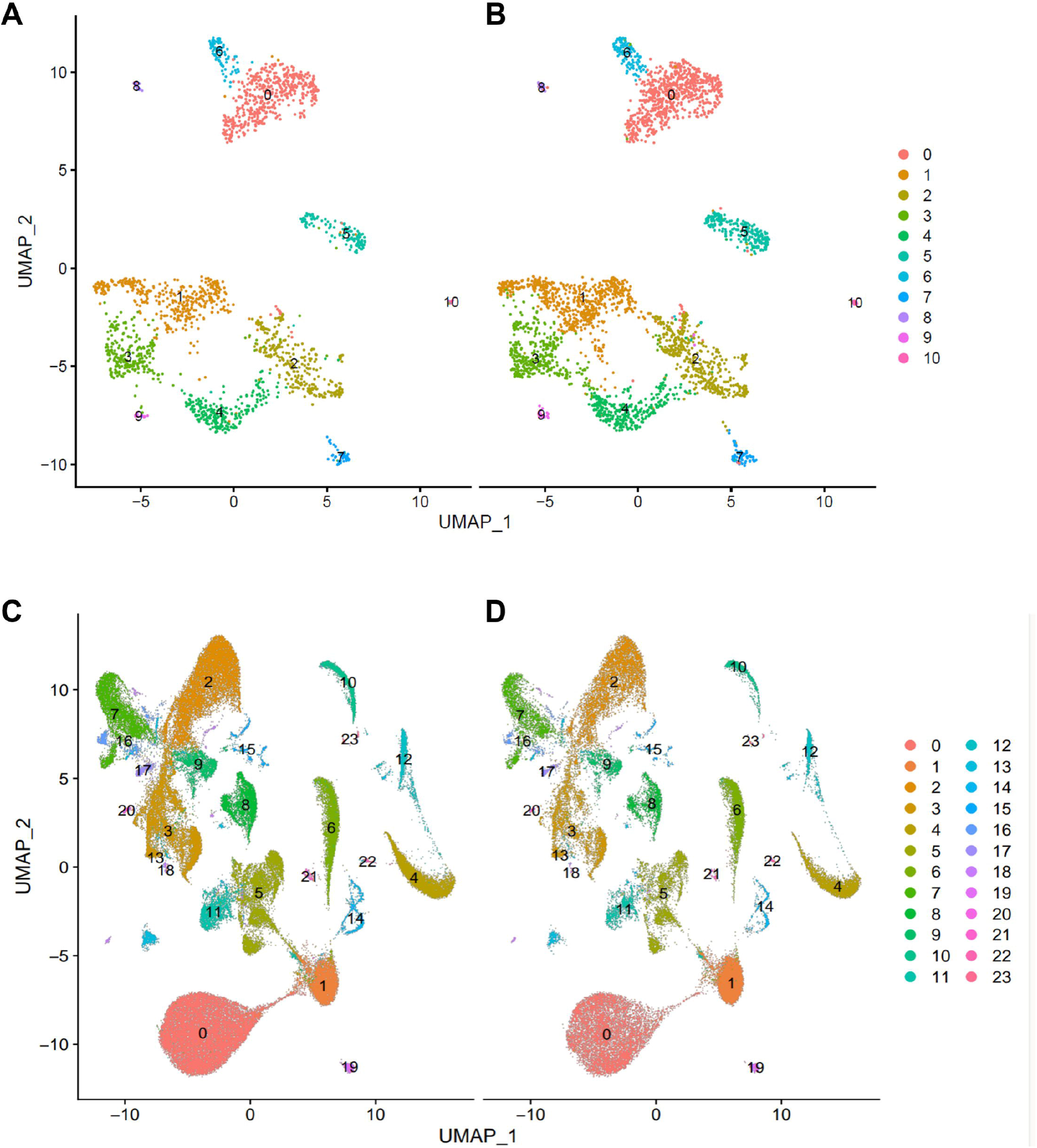
UMAP projections of hippocampal cell clusters in **(A)** control GM^Low^ and **(B)** GM^High^, and cross-foster **(C)** CF^Low^ and **(D)** CF^High^ mice.

**Figure S7.**
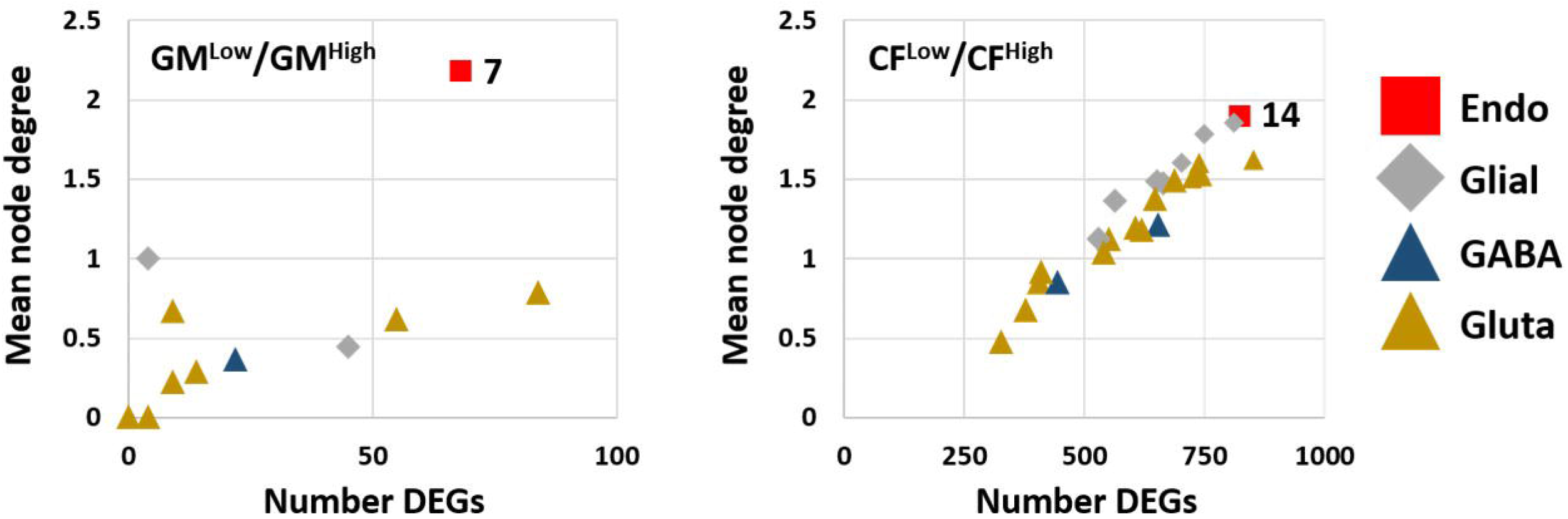
Dot plot correlation between the number of differentially expressed genes (DEGs) and the mean node degree determined by protein interaction analysis of those DEGs, in comparisons of **(A)** GM^Low^ and GM^High^ offspring and in **(B)** CF^Low^ and CF^High^ offspring. Marker shapes denote cell type including endothelial/perivascular cells (Endo, red squares); astrocytes and oligodendrocytes (Glial, grey diamonds); GABAergic neurons (GABA, dark blue triangles); and glutamatergic neurons (Gluta, brown triangles).

**Figure S8.**
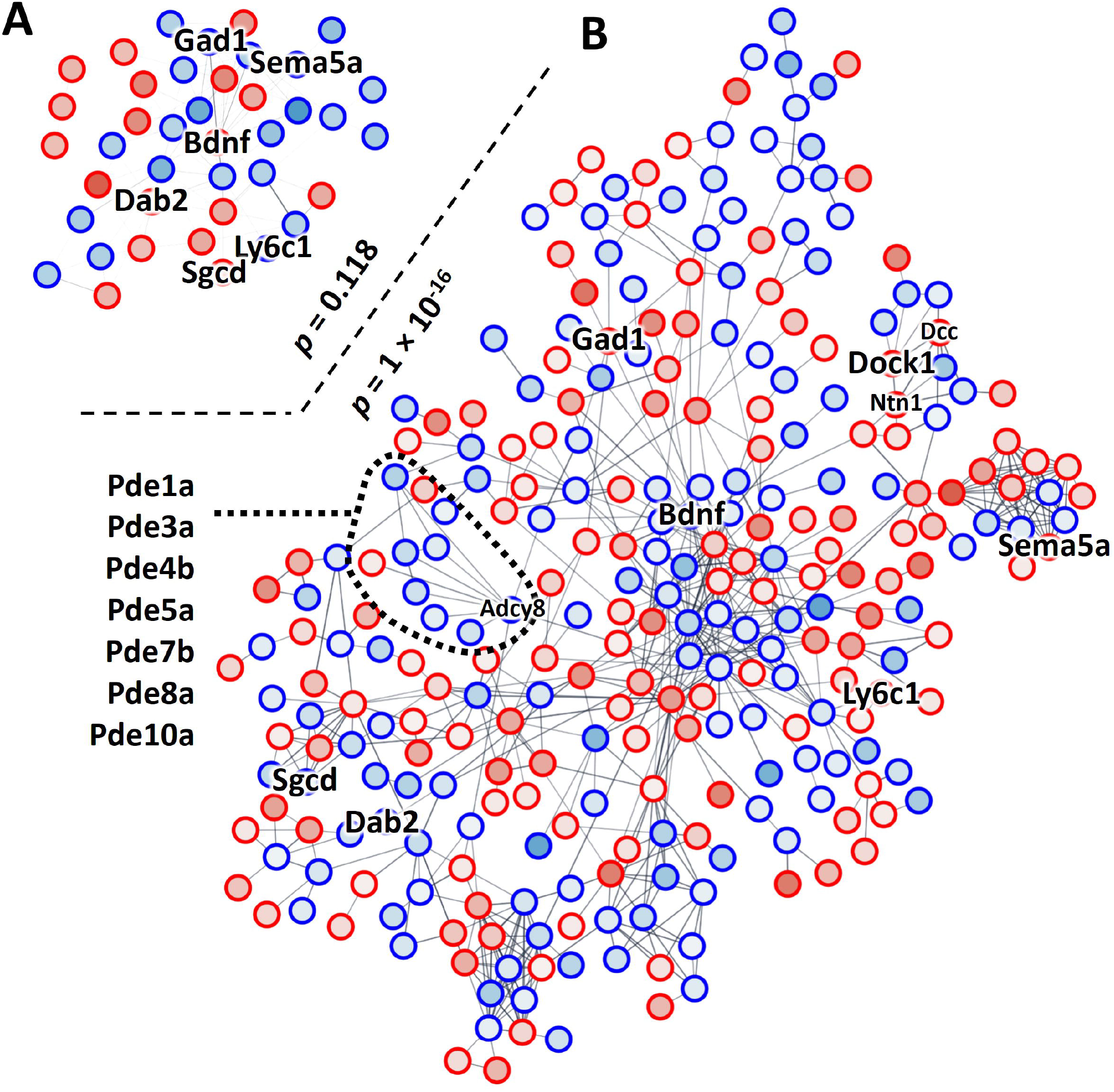
Interaction networks constructed using differentially expressed genes (DEGs) identified in hippocampal level five intratelencephalon (L5 IT)-projecting glutamatergic neurons from **(A)** control and **(B)** cross-fostered mice. Labeled nodes include DEGs identified in both comparisons and showing a pattern of fetal imprinting, and genes that are also differentially methylated (*Dock1*), or closely related (PDE cluster, *Adcy8*, *Dcc*, *Ntn1*).

**Figure S9.**
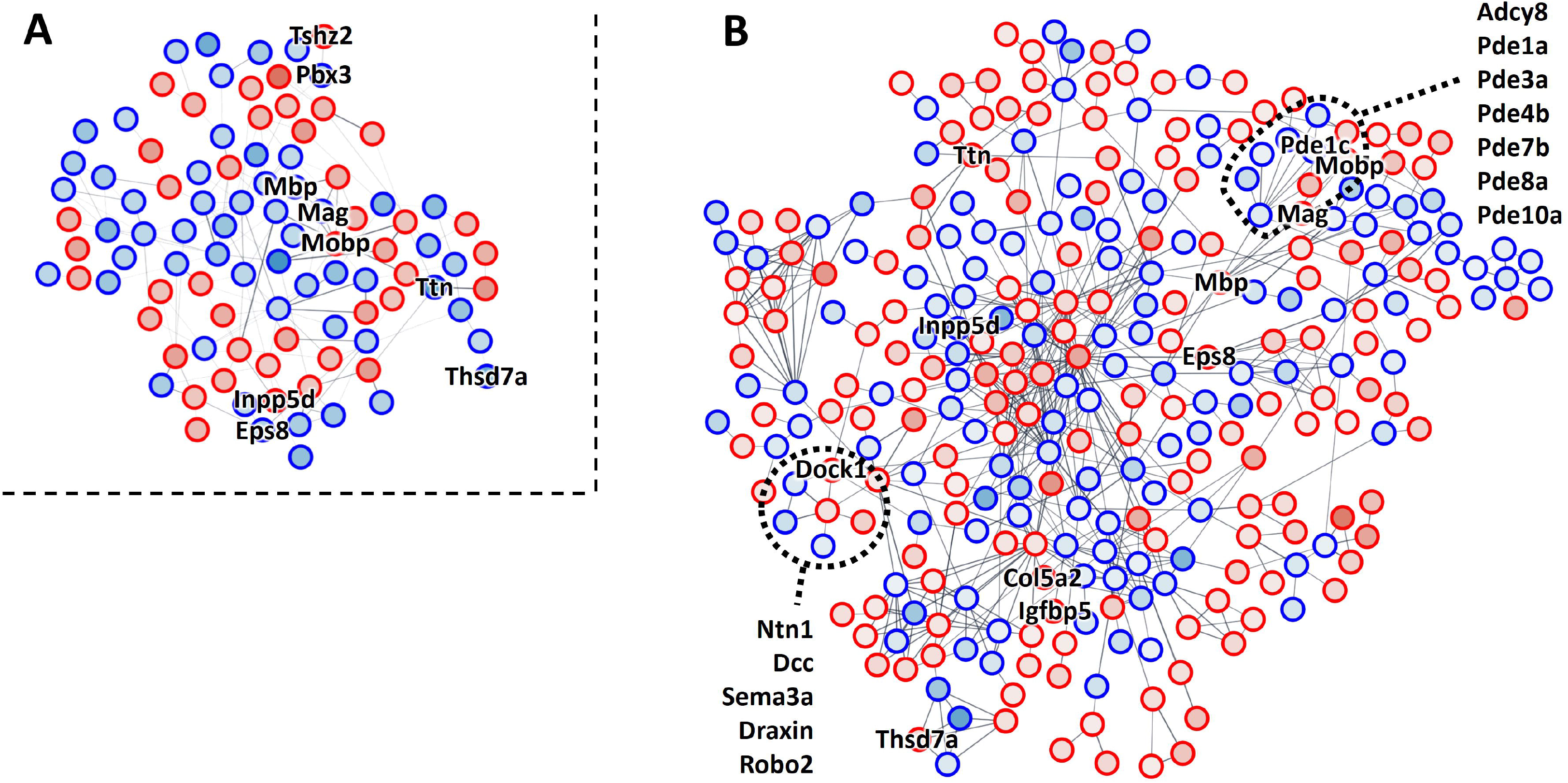
Interaction networks constructed using differentially expressed genes (DEGs) identified in hippocampal astrocyte subsets from **(A)** control and **(B)** cross-fostered mice. Labeled nodes include DEGs identified in both comparisons and showing a pattern of fetal imprinting, and genes that are also differentially methylated (*Dock1*), or closely related (PDE cluster, *Adcy8*, *Dcc*, *Ntn1*).

**Table S1.** Primer pairs used for qRT-PCR analysis.

**Table S2.** Relative abundance of family taxa harbored by either GM^Low^ or GM^High^.

**Table S3.** Fecal metabolite concentrations from GM^Low^ and GM^High^ mice.

**Table S4.** Serum metabolite concentrations from GM^Low^ and GM^High^ mice.

**Table S5.** Relative abundance of identified genera in GM^Low^ and GM^High^ mice.

**Table S6.** Comparison of identified bacterial genera abundance to statistically significant fecal metabolite concentrations between GM^Low^ and GM^High^ mice.

**Table S7.** Differentially methylated regions identified within the genomes of female GM^Low^ and GM^High^ mouse hippocampi.

**Table S8.** Network STRING analysis results utilizing the 196 DMRs identified in the methylome analysis

**Table S9.** Differentially expressed genes identified within the hippocampal cells of GM^Low^ and GM^High^ control male and female mice.

**Table S10.** Differentially expressed genes identified within the hippocampal cells of CF^Low^ and CF^High^ male and female mice.

